# Multimodal foundation model predicts zero-shot functional perturbations and cell fate dynamics

**DOI:** 10.1101/2024.12.19.629561

**Authors:** Zikun Yang, Xueying Fan, Meng Lan, Xubo Tang, Zhijie Zheng, Binghao Liu, Yue You, Luyi Tian, George Church, Xiaodong Liu, Fei Gu

## Abstract

Deciphering cell type-specific perturbation effects on genes and cellular states demands considerable experimental resources. To overcome this challenge, we propose Perturbation Transformer (PertFormer), a foundation model to uncover functional and regulatory mechanisms through interpretable in silico perturbation process. PertFormer comprises 3 billion parameters pretrained on both bulk and single-cell datasets covering 9 types of multiomics (55 billion bp bulk multiomics and the largest-to-date 1.5 billion paired multiomic samples from 1 million single cells), capturing regulation of 300 kb around every genic region. PertFormer enables zero-shot prediction of functional perturbations and cell fate dynamics, outperforming existing methods by 20.9%-480.2%. PertFormer identifies novel tumor treatment targets, validated experimentally. The generalization capabilities of PertFormer have the potential to accelerate the discovery of biological and clinical targets.

---

Regulation of gene expression (*1*) exhibits significant specificity to both cell types and developmental stages (*2*) due to the intricate interplay of transcription factors (*3*) (TFs), chromatin accessibility (*4*), and histone modifications (*5*). To address these questions, substantial efforts have been undertaken: The ENCODE (*6*) project has conducted a large number of epigenetic and transcriptomic experiments on bulk samples, ranging from tissues to cell lines. 10x Genomics (*7, 8*) has sequenced paired gene expression and chromatin accessibility profiles at single-cell or nucleus resolution in a variety of tissues. Although the former initiatives provide a wealth of epigenetic data, they fall short in capturing cellular heterogeneity (*9*). The latter lacks comprehensive multiomic data, as it is challenging to capture such data simultaneously within a single cell (*10*).

Recent advances leverage machine learning, or large language models (*11*) (LLMs) with transformer-based architectures (*12*) to model transcriptional regulation at multiple scales: (a) DNA-related methods (e.g., DeepSEA (*13*), Basenji2 (*14*), BPNet (*15*), scBasset (*16*), EpiGePT (*17*), Enformer (*18*), Borzoi (*19*)), have demonstrated promising results in predicting gene expression or various epigenomic signals from long DNA sequence. (b) Transcriptomics-related methods (e.g., scVI (*20*), scGPT (*21*), scFoundation (*22*), Geneformer (*23*)), showed strong capabilities to extract biologically meaningful representations from high-dimensional transcriptomic data. (c) Multiomics integrated models (e.g., TotalVI (*24*), GLUE (*25*), DRAGON (*26*), EpiBERT (*27*), GET (*28*)), demonstrate strengths in predictions with integrated multimodal information.

Deciphering the molecular determinants of cellular identity and function (*29*), such as master regulators (*30*), provides researchers with powerful tools to precisely engineer cell fate (*31*). By targeting these regulators, desired cell state transitions can be designed (e.g., iPSC reprogramming (*32*), therapeutic treatments (*33*)). Accurate cellular perturbation models are required to evaluate the targets. However, current computational approaches for modeling perturbations on cellular dynamics face fundamental limitations. DNA models elucidate perturbations by sequence mutations, but are not able to simulate epigenetic modifications or targeted gene perturbations. For most transcriptomic models in zero-shot perturbation tasks, their architecture and pretraining strategies are difficult to generate detectable changes between perturbed and wild type outputs, as only a few genes (<0.1%) are modified. Most multiomic-integrated models merely include motif information and open chromatin data, neglecting the long sequence feature as well as chromatin features of histone modifications, TF bindings, which are unavailable in single-cell data.

Moreover, to make predictions on perturbations, most existing models require fine-tuning on high-quality labeled perturbation data (*34, 35*), which is a limited resource due to biological and technical constraints. Existing experimental methods like perturb-seq and CRIPSRi screening only generate the perturbations of limited genes in common cell types, while critical scenarios (e.g., disease microenvironment, multi-genic perturbations, cell state transitions) remain data poor. Consequently, current approaches cannot predict unobserved perturbations without data for fine-tuning, yet biologists lack computational guidance to prioritize targets for perturbation which costs additional experiments.

To overcome the problems of the integration of bulk and single-cell multimodal and elimination of the fine-tuned step in perturbation and cell state predictions, we introduce PertFormer, a 3-billion-parameter foundational model pretrained on both bulk and single-cell datasets covering 9 types of multiomics. The dataset consists of 55 billion bp bulk multiomics and the largest-to-date 1.5 billion paired scRNA and scATAC samples from 1 million single cells, covering regulation of 300 kb around genic region. PertFormer accurately predicted a broad spectrum of tasks through zero-shot (*36*), including the inference of master regulators and GRNs, the predictions of perturbations on transcriptional regulation (*37, 38, 39*). Without the need of labeled data for fine-tuning, PertFormer has demonstrated high accuracy in perturbation predictions from cellular state transitions (*40*) to disease treatments (*41*). Such zero-shot capabilities enable us to discover novel treatment targets in tumor cells, which were further validated. We present all the modules and applications of PertFormer in a comprehensive online website (https://pertformer.ibreed.cn).

## Results

### PertFormer architecture

PertFormer is an attention-based (*12, 42*) multimodal architecture (*43, 44*) pretrained on large-scale multiomic data to uncover functional and regulatory mechanisms through interpretable *in silico* perturbation process in a zero-shot manner (**Fig. 1**). It comprises two principal modules: (1) PertFormer-Elementary to extract high-dimensional features from multiomic data of short-range genomic element (*45*). This module is composed of three layers: the Multiomics embedding (*46*) layer integrates the DNA sequence along with any combinations of multiomic signals as input; the Transformer-1 Block extracts characteristics of each 128-bp bin of the input; and the Transformer-2 Block generates multiomic representation of genomic element from the *R*_[_*_CLS_*_]_ (**Materials and methods**) of all bins (*47*). (2) PertFormer-Regulatory to delineate long-range genomic regulatory patterns (*48*) of a gene. This module targets gene individually, integrated with multiple 1024-bp-sized genomic elements within 300,000-bp flanking the transcription start site (TSS). The multiomic attribute of each element is initially drawn by PertFormer-Elementary, then integrated by attention pooling (*49*). Positional information of the genomic elements is signified through a unique TSS-distance embedding, replacing the traditional position embedding. Coupled with the cross-attention mechanism in the Transformer Block, PertFormer-Regulatory capably simulates the complex long-range interactions among these genomic elements. Overall, PertFormer has over 3 billion neural network parameters (**Fig. S1B**).

**Fig. 1:**
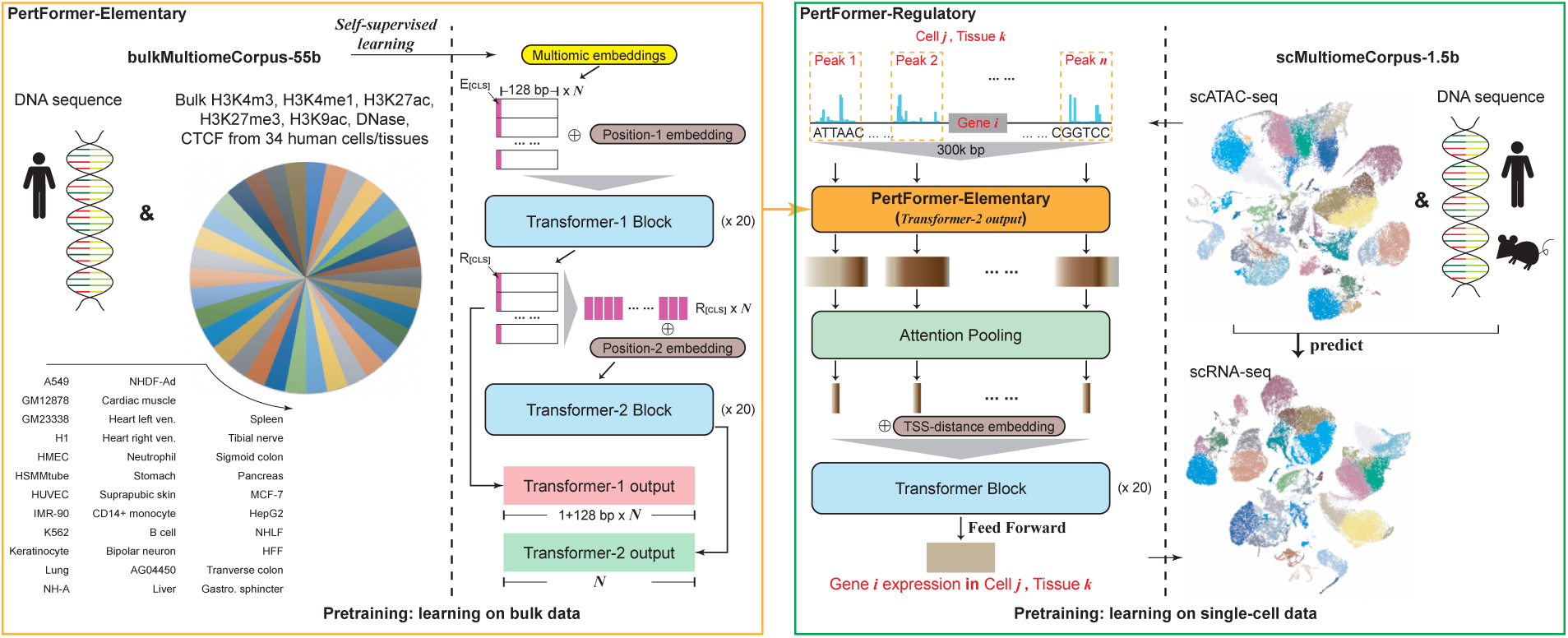
The architecture and pretraining process of the PertFormer model. PertFormer consists of two main modules: PertFormer-Elementary (left) and PertFormer-Regulatory (right). PertFormer-Elementary is designed to process short-range genomic element. The Multiomic embeddings integrates the multimodal information of DNA sequence along with any combinations of epigenetic markers. Two parts of stacked Transformers (Transformer-1, 2) process the input in segmented bins and then merge these bins into an overall multiomic feature. PertFormer-Regulatory employs cross-attention mechanisms among multiple genomic elements within a 300,000-bp range around each gene, using PertFormer-Elementary to process each genomic element, and thereby creating its long-range regulatory features. Two comprehensive databases, bulkMultiomeCorpus-55b and scMultiomeCorpus-1.5b, were proposed for PertFormer pretraining. The bulkMultiomeCorpus-55b contains various gennomic elements of DNA sequence along with 7 types of epigenetic markers across 34 bulk human cell lines/tissues, with a total length of over 55 billion bp. The scMultiomeCorpus-1.5b is an extensive single-cell database built from paired scATAC and scRNA data in 1 million single cells, comprising regulatory relations across 1.5 billion of the 300kb genic regions. PertFormer-Elementary was pretrained by self-supervised learning in bulkMultiomeCorpus-55b. PertFormer-Regulatory was pretrained on scMultiomeCorpus-1.5b, leveraging DNA sequences and open chromatin profiles to predict the expression levels of massive genes in massive single cells. PertFormer-Regulatory was shared across all genes and all cells

To pretrain PertFormer, we proposed two comprehensive databases, bulkMultiomeCorpus-55b and scMultiomeCorpus-1.5b, derived from publicly available bulk and single-cell data. The bulkMultiomeCorpus-55b was built from the ENCODE bulk epigenetic profiles, focusing on key markers of H3K4me3, H3K4me1, H3K27ac, H3K27em3, H3K9ac, DNase, and CTCF across 34 human cell lines and tissues (**Data S1**), together with DNA sequence. This corpus encapsulates millions of genomic regions cumulatively that span beyond 55 billion bp. The scMultiomeCorpus-1.5b, on the other hand, was curated by an extensive collection of scMultiome data provided by the 10x Genomics platform (**Data S1**), mapping out the largest-to-date 1.5 billion paired scRNA and scATAC associations from 1 million single cells.

The pretraining of PertFormer-Elementary was conducted through self-supervised learning (*50*), utilizing the bulkMultiomeCorpus-55b (**Fig. S1A**). Briefly, the multiomic input was randomly partitioned into Selected and Preserved groups. PertFormer-Elementary was trained to predict the Preserved group from the Selected group. Concurrently, Transformer-1 Block optimized its output in executing masked and reconstruction tasks on the Selected group. After pretraining of PertFormer-Elementary and freezing its weights, we advanced to the pretraining of PertFormer-Regulatory. In diverse single-cell contexts, this stage concentrated on predicting the expression levels of vast genes based on DNA and chromatin accessibility profiles from all consensus scATAC peaks within 300,000-bp flanking the TSS of each gene, leveraging the scMultiomeCorpus-1.5b dataset. Specifically, the parameters of PertFormer-Regulatory were shared across all genes.

In the following results, no task-specific data was used to either fine-tune any parameters in the pretrained PertFormer, or build any additional downstream models. Such zero-shot capabilities of PertFormer were demonstrated across multiple applications in sequence-level, chromatin-level, gene-level, and cell-level scenarios.

### Zero-shot predictions of master regulators and GRNs

The PertFormer’s attention mechanism is able to detect key genomic elements that regulate gene expression. To interpret functional GRNs, two key metrics derived from this approach are proposed: the Attention Score, quantifying the regulatory importance of genomic elements on a specific gene, and the Attention Feature, encapsulating the overall regulatory profile among genes (**Materials and methods**).

In B cells of PBMCs featured in scMultiomeCorpus-1.5b, PertFormer-Regulatory’s Attention Score (*51*) yielded mean Spearman correlation coefficient (SCC) (*52*) of 0.810 with ChIP-seq signals of B cell’s master regulators, PAX5, EBF1, and POU2F2 (*53*) (**Fig. 2A**, pretrained cells). An example of the 300,000-bp flanking the TSS of the RPL28 gene was displayed, where regions with high Attention Score were closely aligned with the intense ChIP-seq signal of EBF1, PAX5, and POU2F2, and vice versa (**Fig. 2B**). Furthermore, the Attention Score’s predictive power can be generalized to cells not present in scMultiomeCorpus-1.5b, as demonstrated by a mean SCC of 0.783 with ChIP-seq signal of SP1 and FOSL1 (*54*) in colorectal cancer (CRC) tumor cells (**Fig. 2A**, Unseen cells). For comparison, SCC for pseudobulk scATAC-seq signal was 0.582. Thus, the Attention Score of PertFormer model can pinpoint functional elements that correlate with the binding of master regulator (*55*).

**Fig. 2:**
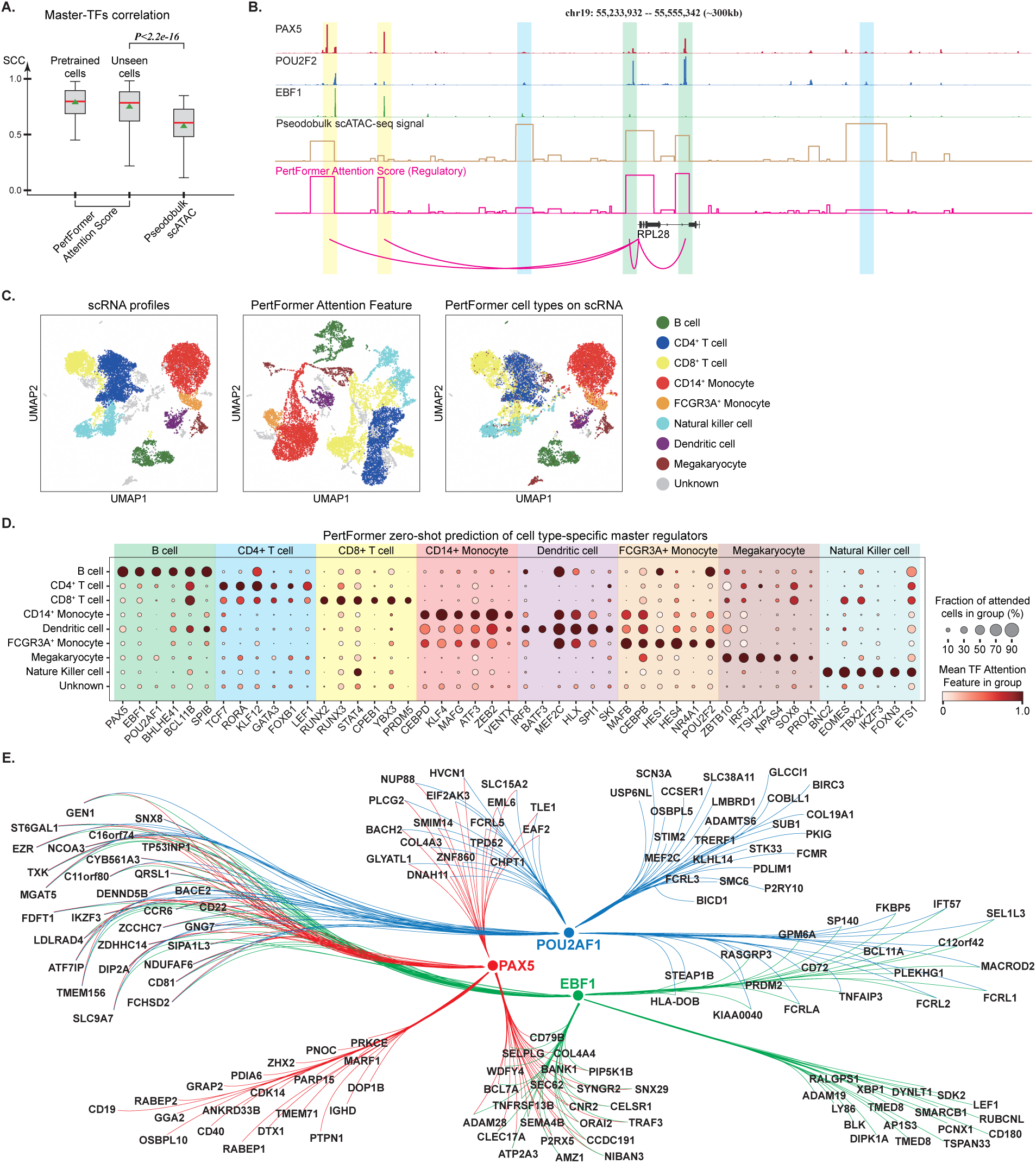
Zero-shot predictions of master regulators and GRNs. **A.** Distribution of SCC between PertFormer Attention Score in each context (pretrained-cells, all genes in B cells of PBMCs; Unseen cells, all genes in tumor cells of CRC) versus master-TFs’ ChIP-seq signals. Each value was an SCC for a specific gene in a specific context. Pseudobulk scATAC was used as control group. PertFormer Attention Score exhibited higher correlations with master-TF signals and can be generalized across various genes and cells (*P* < 2.2 × 10^−16^ one-sided Mann-Whitney U test; centre lines, median; triangles, mean; box limits, upper and lower quartiles; whiskers, 1.5× interquartile range). **B.** PAX5, POU2F2, and EBF1 TF ChIP-seq signals, pseudobulk scATAC profiles, together with PertFormer Attention Score of 300,000-bp flanking the RPL28 gene in B cells of PBMC. PertFormer’s predicted region-gene links were shown as arcs. Strong master-TF bindings were found in distal (yellow) and proximal (green) PertFormer-highly-attended regions for this gene. In contrast, regions with high chromatin accessibility (blue) demonstrated low TF signals, and showed low PertFormer Attention Score. **C.** UMAP of PBMC single cells based on experimental scRNA profiles (left) and PertFormer Attention Feature (middle). Cell types annotated by Attention Feature were marked on the UMAP of experimental scRNA (right). Cell type annotation results between scRNA and PertFormer showed a high consistency of 0.809 (right). **D.** Cell type-specific master regulators predicted by PertFormer Attention Feature. Mean Attention Features were shown by color scale, while fraction of cells were shown by size scale. Many well-known master regulators of each cell type were recovered by PertFormer. **E.** PertFormer-predicted GRNs regulated by PAX5, EBF1 and POU2F2 in B cells of PBMCs. These 3 TFs formed 7 distinct groups of regulatory combinations.

Leveraging PertFormer attention mechanisms, we proposed a workflow to infer functional GRNs. Taking PBMCs as an illustrative case for included-cell prediction, we first applied dimensionality reduction (*56*) on PertFormer Attention Feature to cluster the single cells on Uniform Manifold Approximation and Projection (*57*) (UMAP). The results not only showed clearly separated groups for different cell types, but also matched closely (with a concordance of 80.9%) with cell type annotations derived from scRNA profiling (**Fig. 2C**).

We subsequently identified TFs with distinct Attention Feature (**Materials and methods**) across each cell types (**Fig. 2D**). PertFormer *in silico* explorations illuminated critical cell type-specific TFs. Many well-known master regulators (*58*), including PAX5, EBF1, POU2F2, and SPIB of B cells (*59*), TCF7, LEF1, RUNX2, and STAT4 of T cells (*60*), CEBPD, NR4A1, CEBPB, and HES4 of monocytes (*61*), IRF8 and ST18 of dendritic cells (*62*), as well as BNC2, EOMES, and TBX21 of natural killer (NK) cells (*63*), were recovered by PertFormer Attention Feature. These predicted TFs showed significant differential expressions (**Fig. S3**). As control, master regulators cannot be predicted using scATAC signals (**Fig. S3**).

Next, through motif analysis (*64*) on highly-attended functional regions, PertFormer mapped the predicted master regulators to specific genes to construct GRNs. For 150 highly-variable genes in B cells of PBMCs, we scanned for PAX5, POU2F2 and EBF1 motifs in regions with significant Attention Score. Motif enrichment across these genes facilitated their categorization into seven distinct groups (**Fig. S4A**), laying out the inferred GRNs via PertFormer (**Fig. 2E**). This regulatory schema yielded 46.7% consistency with SCENIC+’s predictions, and recaptured 67.4% of SCENIC+’s TF-to-gene relationships (*65*).

Notably, PertFormer’s utility in inferring GRNs relies on solely DNA sequences and scATAC-seq profiles, obviating the need for paired scMultiome datasets or fine-tuning to produce predictions in novel cell types outside of scMultiomeCorpus-1.5b. This was supported by an unseen-cell application of colorectal cancer (CRC) tumor tissue (**Fig. S2, S4B**). PertFormer precisely demarcated tumor from normal cell types (80.2% conformity to scRNA-derived annotations), and identified prominent CRC tumor-specific master regulators, such as CDX2, CDX1, FOSL2, HOXB9, HOXA13 and ELF3 (*54*). Moreover, PertFormer unveiled GRNs within CRC tumor cells.

### Zero-shot predictions of perturbations on transcriptional regulation

Utilizing experimental methods such as CRISPR interference (*66*) (CRISPRi) to delineate the regulatory effects of genomic regions (*67,68,69*) is commonly time-consuming and expensive (*70*). The consequent scarcity of experimentally validated perturbation responses substantially hampers computational models for *in silico* predictions (*71*). We herein presented PertFormer as a pioneering zero-shot approach to pinpoint functional targets and simulate their knockout (KO) and knock-in (KI) effects.

To prioritize functional enhancer-gene pairs (EG-pairs), bulk ATAC profiles were utilized to identify enhancer candidates and compute their Attention Scores. Specifically, for each gene, enhancer candidates were defined as chromatin-accessible peaks spanning 300,000-bp around the TSS and ranked by their Attention Scores on this gene. For the K562 CRISPRi-FlowFISH dataset (*72,73,74*), PertFormer demonstrated higher rank of Attention Scores for validated true EG-pairs than false ones (**Fig. 3A**, left). For control group, the ATAC signals did not show a substantial difference. In comparison with the best existing methods, PertFormer yielded an average of 60.3% better performance in auPRCs to predict EG-pairs with different regulatory distances (from 11.2% to 160.3%, **Fig. 3A**, right). The performance of PertFormer was further evaluated using EG pairs collected from the VISTA mouse enhancer database (*75, 76*), an average of 116.4% improvements in auPRC were observed across different EG-pair distances (from 10.7% to 350.4%, **Fig. 3B**). As an illustrative case of PQBP1 gene in K562 (**Fig. 3C**), PertFormer allocated strong Attention Scores to the four CRISPRi-validated true enhancers (green), whereas it assigned lesser Attention Scores to CRISPRi-affirmed false enhancers (gray).

**Fig. 3:**
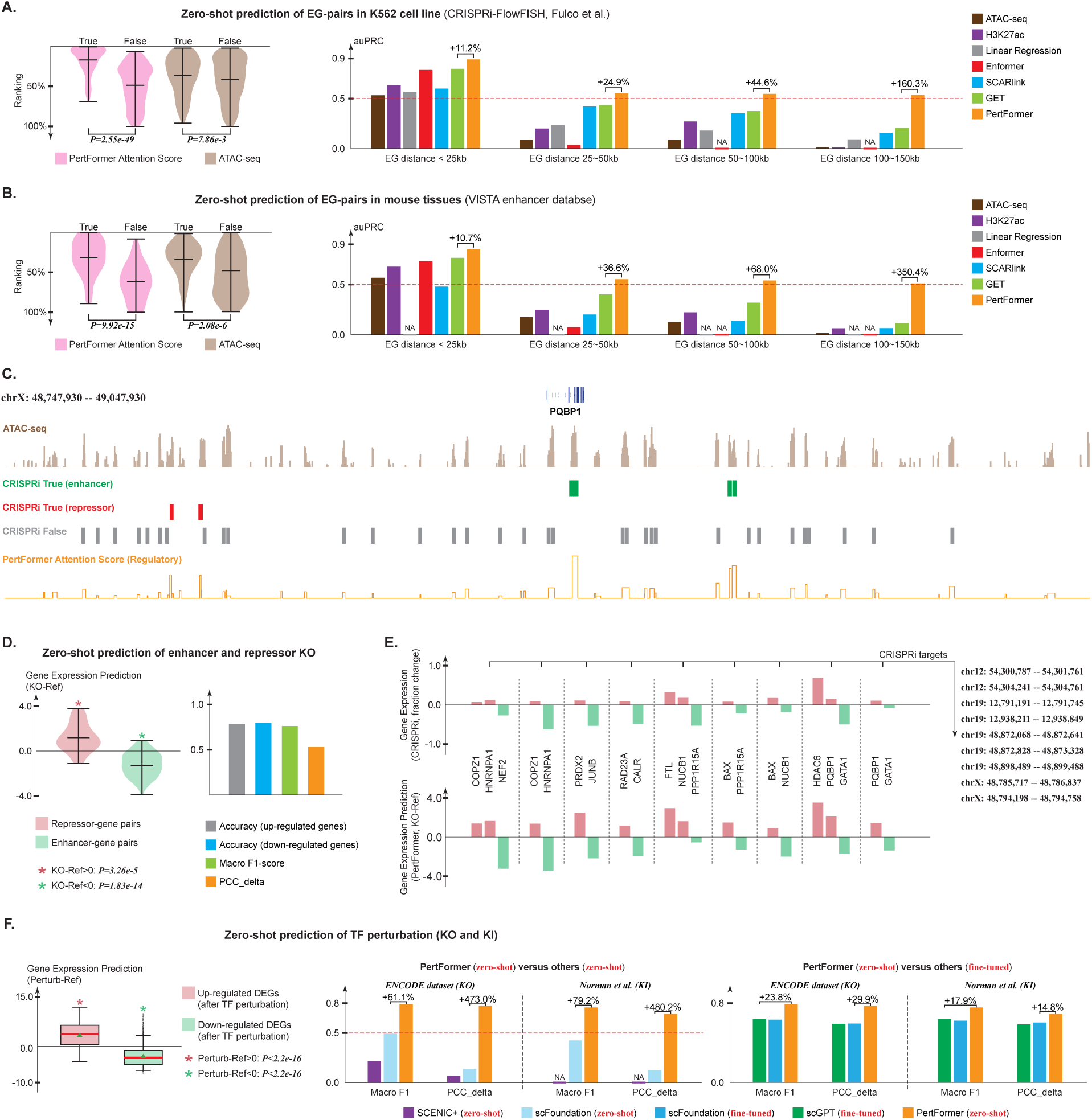
Zero-shot predictions of perturbations on transcriptional regulation. **A.** Violin plots showed distribution of ranking of CRISPRi-validated true and false EG-pairs based on PertFormer Attention Score or ATAC signal. Attention Scores were generated from PertFormer-Regulatory Transformer Block using bulk ATAC data. PertFormer Attention Scores on true EG-pairs were significantly higher than false ones (P-values, one-sided Mann-Whitney U test). Bar plots showed the auPRCs of EG-pairs prediction. Across different regulatory distances, PertFormer demonstrated 11.2%-160.3% better performance in comparison with the best existing methods. **B.** Ranking distributions and auPRCs for true and false EG-pairs in mouse VISTA enhancer database. PertFormer was 10.7% to 350.4% better than the best existing methods to predict EG-pairs across different regulatory distances. **C.** Bulk ATAC-seq profiles, PertFormer Attention Score, and CRISPRi-FlowFISH experimental results in 300,000-bp region flanking the TSS of PQBP1 gene in K562 cell line. Locations of validated true or false enhancers that regulate PQBP1 were respectively marked in green or gray. PertFormer exhibited high Attention Scores on the 4 validated true enhancers and low attentions on the false ones. ATAC signals were consistently high across all true and false enhancers. Two CRISPRi targets validated as repressive effects were marked in red. PertFormer showed high attentions on these two repressors. **D.** Violin plots showed distribution of PertFormer-predicted gene expression changes (KO-Ref) after *in silico* KO at enhancer or repressor loci. *In silico* KO at repressor loci (red) showed up-regulated gene expressions, and vice versa for enhancers (green). P-values were acquired by one-sided Mann-Whitney U test to see if the predicted KO-Ref were significantly larger or smaller than 0, respectively. Centre lines represented the mean values. Bar plots showed the performance to distinguish enhancers and repressors. Accuracy and macro-F1 were computed by setting 0 as threshold to reflect the predicted changing direction of gene expression. PCC_delta was measured between the predicted and experimental gene expression changes. **E.** Fraction changes of gene expression after CRISPRi, and PertFormer-predicted gene expression changes (KO-Ref), for 9 unique CRISPRi targets. Each of those targets has bivalent regulatory effects on different genes, which was correctly modeled by PertFormer. **F.** Box plot showed distribution of PertFormer-predicted gene expression changes after *in silico* TF perturbation (KO and KI) in K562 cell line (P-values were computed by one-sided Mann-Whitney U test to evaluate if the predicted KO-Ref were significantly larger or smaller than 0, respectively for the up- or down-regulated DEGs; centre lines, median; triangles, mean; box limits, upper and lower quartiles; whiskers, 1.5× interquartile range). Bar plots showed the performance to model the TF perturbations. Macro-F1 was computed by setting 0 as threshold to reflect the predicted changing directions of gene expression. PCC_delta was measured between the predicted and experimental gene expression changes. In zero-shot mode, PertFormer showed 61.1% (KO) and 79.2% (KI) better macro F1, as well as 473.0% (KO) and 480.2% (KI) better PCC_delta than the best existing methods. Zero-shot PertFormer was still better than other fine-tuned methods.

The K562 CRISPRi-FlowFISH study also validated genomic elements with significant repressive regulatory effects on gene expression. Notably, 79.3% of these elements were ranked in the top quartile by PertFormer’s Attention Scores among all CRISPRi targets (for example, in **Fig. 3C**, the two CRISPRi-validated repressors also received strong attention from PertFormer). This inspired us to further explore PertFormer’s capacity to infer the regulatory effects of both enhancers and repressors (*77*). We introduced *in silico* KO simulation at genomic loci of enhancers or repressors by erasing their chromatin accessibility signals, and assessed their influence on gene expression. We performed this *in silico* KO on each of the CRISPRi-validated true EG and repressor gene (RG)-pairs in K562 cell line. PertFormer’s responses to gene expression were in close agreement with CRISPRi experiments, as 78.2% of genes that were experimentally observed to be up-regulated and 79.5% of those observed to be down-regulated after CRISPRi were correctly predicted (**Fig. 3D**), resulting in a macro-F1 score of 0.759 (**Fig. 3D**, right). In addition, the predicted changes in gene expression showed 0.525 in Pearson correlation coefficient delta (*78*) (PCC_delta) with CRISPRi results (**Fig. 3D**, right). Notably, 13 of those CRISPRi targets that had repressive effects on certain genes were also validated as enhancers for other genes. For 69.2% of these targets, PertFormer correctly predicted their regulatory effects (**Fig. 3E**).

To evaluate TF perturbation predictions, the ENCODE TF perturbation datasets (*6*) (TF KO) and the Norman perturb-seq datasets (*79*) (TF KI) were used. In perturbation simulations, *in silico* TF KO was modeled by removing ATAC signals in regions enriched with the motifs of the corresponding TF, while *in silico* TF KI was conducted by insertion of ATAC signals into genomic loci identified with the corresponding TF motif (**Materials and methods**). PertFormer predicted changing directions of differentially expressed genes (*80*) (DEGs) concordant to experimental perturbations (**Fig. 3F**, left), achieving macro-F1 of 0.789 for KO, and 0.754 for KI (**Fig. 3F**, middle). Besides, PertFormer exhibited 0.768 (KO) and 0.690 (KI) in PCC_delta to model the changed values in genome-wide gene expressions (**Fig. 3F**, middle). In zero-shot mode, existing models showed low predictive capability (**Fig. 3F**, middle, macro-F1<0.5, PCC_delta<0.15). Remarkably, compared with existing models that had been fine-tuned with perturb-seq data, the zero-shot PertFormer was still better (**Fig. 3F**, right, 14.8%-29.9% improvements). As control, random loci were used to perform the perturbation test, and showed no predictive capability (**Fig. S5**, **Supplementary Text**), further demonstrating the effectiveness of selecting TF motifs as perturbation loci.

### Zero-shot predictions of cell state transitions

Cell state transitions can be induced by activation of key regulators to produce desired cell types (e.g., higher pluripotency (*81*)). Existing methods face limitations due to the scarcity of fine-tuning data in such scenarios, which can be addressed by PertFormer’s zero-shot capability. Here, we showed PertFormer simulations of the reprogramming effects from somatic cell types to induced pluripotent stem cell (*82*) (iPSC).

We first simulated the effects of iPSC reprogramming in three types of human somatic cells, fibroblasts, cardiomyocytes, and B cells, which were known cell types that can be reprogrammed by the 4 pluripotency TFs (*83*), OCT4, SOX2, KLF4, and c-MYC (OSKM) (*84, 85, 86*). We applied PertFormer to model the *in silico* KI effects of OSKM by inducing the highest ATAC peak flanking the 300,000-bp of the TSS of targeted gene to the loci with OSKM motifs (**Materials and methods**). To evaluate the performance of gene expression prediction, we identified numerous DEGs by bulk RNA-seq between human embryonic stem cell (hESC) and these three types of somatic cells. In *in silico* KI simulation across these somatic cell types by PertFormer, down-regulated DEGs after differentiation experienced reversal and up-regulation effects, and vice versa (**Fig. 4A**). These findings are consistent with the iPSC reprogramming effects (*84*).

**Fig. 4:**
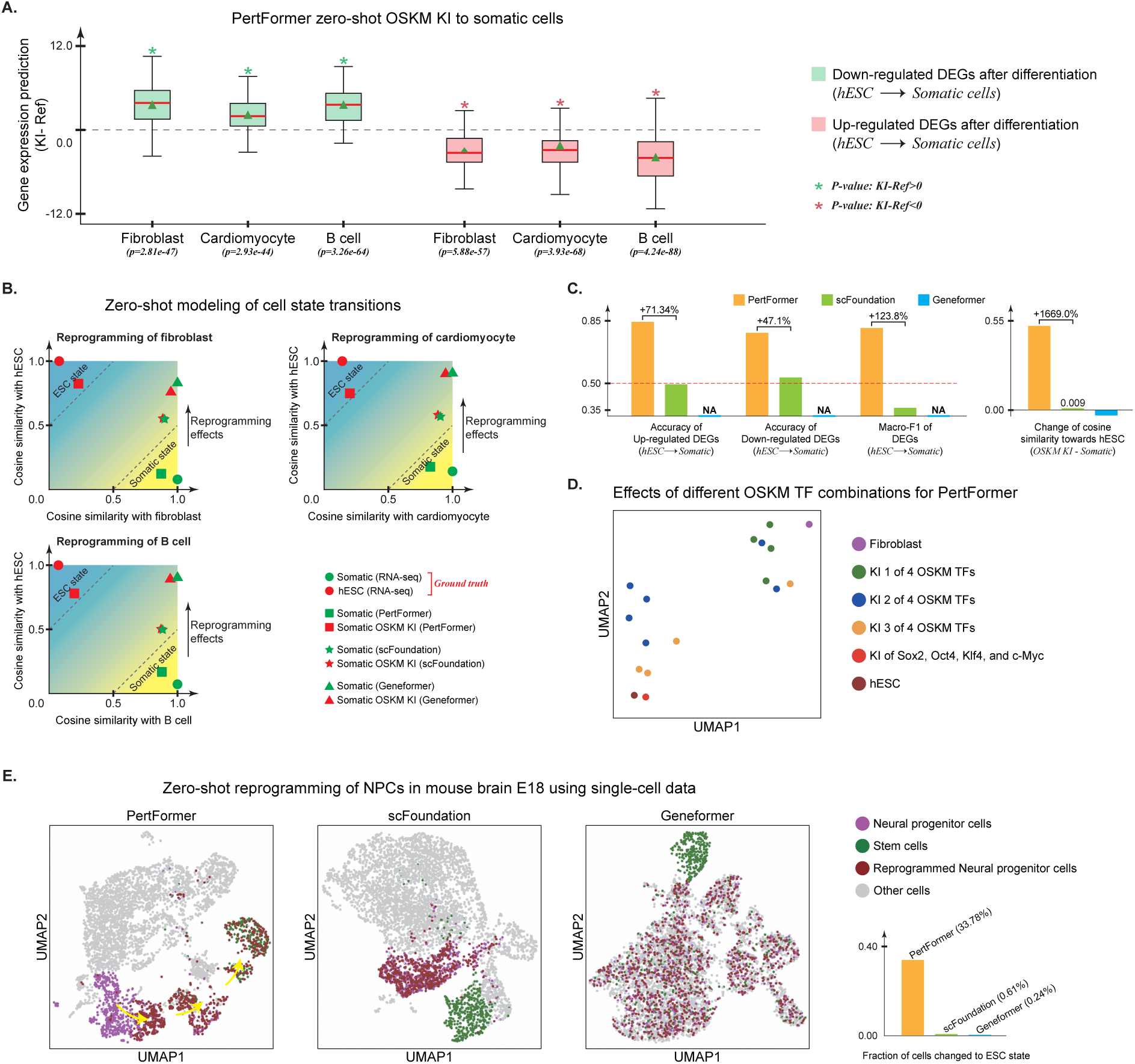
Zero-shot predictions of cell state transitions. **A.** PertFormer OSKM KI to somatic cells. DEGs were identified by RNA-seq from hESC to each somatic cell type. Box plots showed distribution of predicted gene expression changes (KI-Ref) for each DEG in each of the 3 somatic cell types. For down-regulated DEGs after development, KI increased their expressions, and vice versa. P-values, one-sided Mann-Whitney U test; Centre lines, median; triangles, mean; box limits, upper and lower quartiles; whiskers, 1.5× interquartile range; points, outliers. **B.** *In silico* OSKM KI-induced cell state transitions reflected by cosine similarity. After *in silico* OSKM KI, PertFormer is the only model that showed clear cell state transitions from somatic state to ESC state, across all 3 somatic cell types. **C.** Accuracy, macro-F1 and changes of cosine similarity towards hESC, after *in silico* OSKM KI to somatic cells. Existing methods showed low predictive power. **D.** UMAP of fibroblast, hESC, and *in silico* reprogrammed cells with KI of different OSKM TF combinations, based on experimental (hESC and fibroblast) or predicted (reprogrammed cells) gene expression differences with fibroblast. With increased number of OSKM TFs used for KI process, PertFormer-simulated reprogrammed cellular states were closer to hESC. **E.** UMAPs showed NPCs, mESCs, along with OSKM reprogrammed NPCs in mouse embryonic brain E18, based on PertFormer Attention Feature, Geneformer cell embedding, and scFoundation cell embedding using single-cell samples. Only in PertFormer, cell state transition from NPCs to mESCs can be observed after KI. Bar plot showed fraction of cells changed to ESC state after *in silico* reprogramming.

Next, cell state transitions during *in silico* reprogramming were evaluated by cosine similarity between predicted and wild-type gene expression profiles from somatic to iPSC cells (**Fig. 4B**). For cells in somatic state, PertFormer predicted gene expression profiles showed an average of high cosine similarity of 0.877 with the somatic state and a low cosine similarity of 0.215 with the ESC state. After *in silico* OSKM KI by PertFormer, strong reprogramming effects were observed, as the perturbed cell transitioned to ESC-like states (0.747 and 0.234 cosine similarity with hESC and somatic cells, respectively).

Overall, PertFormer-simulated OSKM inductions resulted in an averaged macro-F1 of 0.795 to infer the changing directions of DEGs (**Fig. 4C**, left), as well as the change of cosine similarity of 0.532 to reflect the cell state transitions from somatic cells to hESC (**Fig. 4C**, right). Notably, existing methods showed very limited predictive power (**Fig. 4C**).

In addition, reprogramming effects of different combinations of OSKM were predicted by PertFormer. In bulk samples, we used dimensionality reduction to depict cell state transitions based on the experimental and PertFormer-predicted gene expression changes during reprogramming with different OSKM combinitions (**Materials and Methods**). Reprogrammed fibroblast displayed a progressive reversion toward the hESC state, escalating incrementally with the number of OSKM TFs employed for KI (**Fig. 4D**). A near-complete restoration to the stem cell state was achieved when cells were reprogrammed using all 4 OSKM TFs.

Furthermore, prediction of cell state transitions was also tested in scRNA profiles of mouse embryonic brain E18. Neural progenitor cell (NPC) was used for simulation, as its reprogramming by OSKM has been reported before (*87*). We marked NPCs and mouse embryonic stem cells (mESCs) based on marker gene expressions (**Data S2d**). Using scATAC data, we performed PertFormer *in silico* reprogramming through KI of OSKM on all NPCs. Clear transition of NPC clusters towards the stem cell state was observed, and 33.78% of the reprogrammed NPCs reached mESC state (**Fig. 4E**). In comparison, other approaches exhibited negligible effects (**Fig. 4C**, right).

### Zero-shot predictions of novel treatment targets

Transcriptional regulation on targeted genes contributes to the formation, progression and treatment of human cancer (*88,89*). Here, we applied PertFormer to model therapeutic effects in various cancer types (*89*) in a zero-shot manner.

We first applied PertFormer to predict tumor-specific master regulators across 5 types of primary tumors (*90*). For each tumor type, top 6 predicted TFs were shown and their tumor-promoting (red) or tumor-suppressive (green) effects (**Fig. S6A**) were obtained according to previous studies (**Supplementary Text**). We further investigated whether such regulatory effects can be simulated by PertFormer. For each tumor type, perturbations (KO or KI) of top 1 tumor-promoting and top 1 tumor-suppressive TFs were simulated by PertFormer. The KO or KI perturbation was chosen if the gene expressions level of a TF in tumor cells was higher or lower than that of normal cells (**Fig. S6B**). Compared with original UMAP of normal and tumor cells of the 5 tumor tissues (**Fig. S6C**), PertFormer correctly predicted the direction of cell state transitions of both tumor-promoting and tumor-suppressive TFs (**Fig. S6D**). For example, KO of tumor-promoting TFs (ESR1, MECOM, GATA2, and TP73) resulted in cell state transitions from tumor cells towards closest normal cells (CNCs), KI of tumor-promoting TF AR caused the tumor cells to transit away from CNCs, KO of tumor-suppressive genes (RORA, NR3C2, HNF4A, THAP5, and TCF3) showed tumor cells shifted away from CNCs (**Fig. S6D**). Therefore, PertFormer can identify functional targets across different tissues and predict their promoting or suppressive effects on tumors.

Encouraged by this, we further used PertFormer to search for novel treatment targets in triple-negative breast cancer (TNBC) and ovarian cancer (OV). Top 6 and top 8 master regulators of TNBC and OV were well known from previous studies (**Fig. 5A**, **Supplementary Text**). PertFormer simulated perturbations of these targets showed concordance with their known tumor-promoting or suppressive effects (**Fig. S7A-C**, **Supplementary Text**). Moreover, THAP2 in TNBC, and ZIK1 in OV, were identified as novel targets that have never been investigated before in tumors. As these two targets exhibited higher expression in tumor cells than normal cells (**Fig. S7A**), KO perturbation were performed by PertFormer and cell state transitions towards CNCs were observed (**Fig. 5B**). Hence, our predictions showed that THAP2 and ZIK1 were potential novel tumor-promoting targets for TNBC and OV tumors, respectively.

**Fig. 5:**
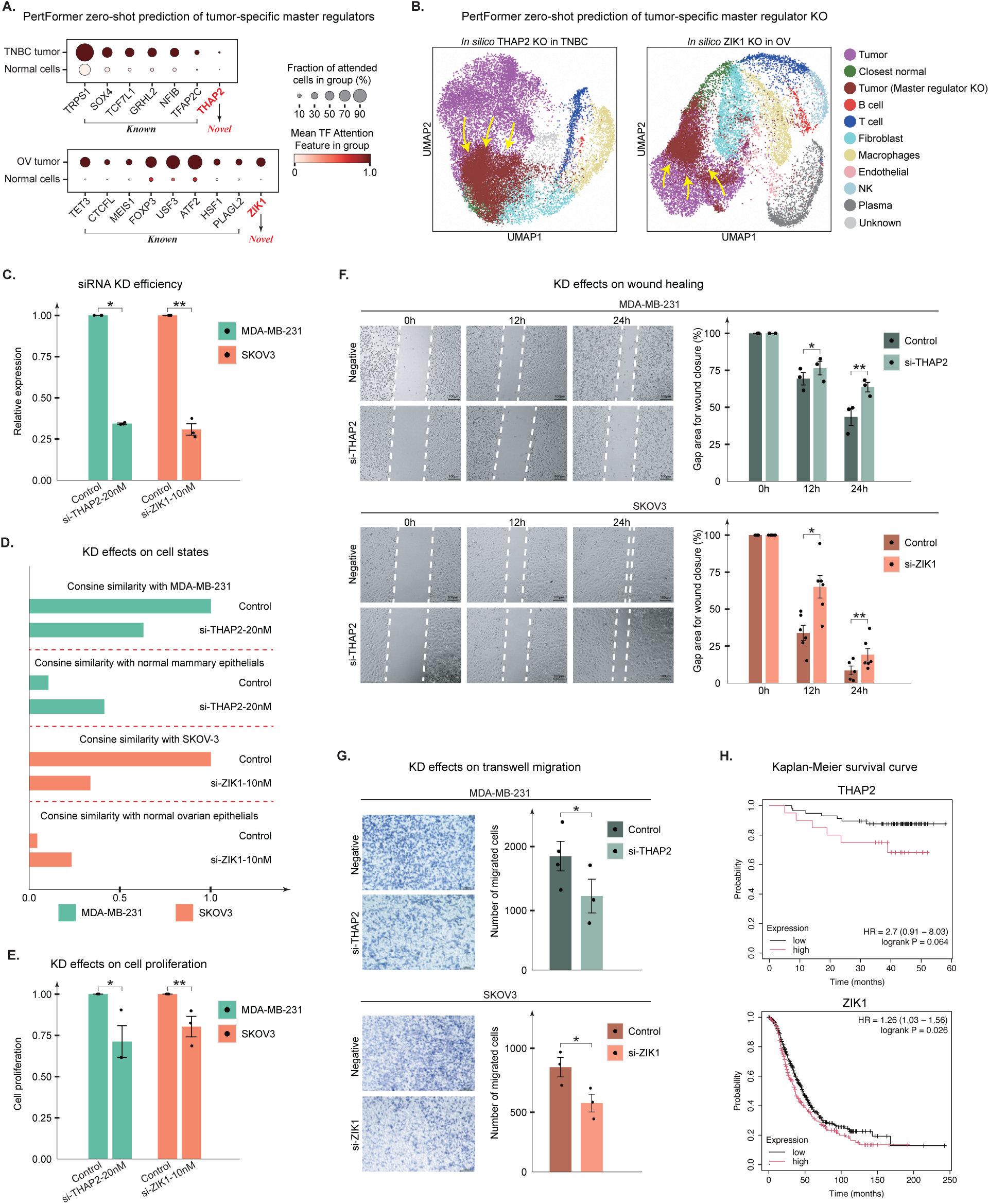
Zero-shot predictions of novel treatment targets. **A.** Master regulator predictions in TNBC and OV, based on PertFormer Attention Feature. In tumor cells, the top 6 TFs for TNBC and the top 8 TFs for OV, were reported by previous studies. THAP2 in TNBC and ZIK1 in OV were identified as novel targets. Mean TF Attentions and fraction of cells were shown by color scale and size scale, respectively. **B.** Predicted perturbation effects of novel targets. KO simulations were performed, as the expressions of the two targets were higher in tumor cells. UMAP showed normal, tumor, and *in silico* TF-KO tumor cells based on PertFormer Attention Feature. Both targets were predicted as tumor-promoting, as the KO cells (brown dots) showed clear cell state transitions towards the CNCs (green dots). **C.** Validation of siRNA KD efficiency for THAP2 in MDA-MB-231 and ZIK1 in SKOV3 using qPCR. siRNA-mediated KD resulted in significantly dropped expression levels (*p=3.57e-5, **p=2.00e-3, t-test). **D.** Changes of gene expressions after siRNA-mediated KD. Bulk RNA-seq profiles were used to measure the gene expressions for KD and control groups in both cancer cell lines. After KD, gene expressions showed decreased cosine similarity with TNBC or OV cell lines, and increased cosine similarity with normal mammary or ovarian epithelial cells. **E.** Cell proliferation after 48 h was measured using the CCK-8 assay following THAP2 KD in MDA-MB-231 and ZIK1 KD in SKOV3. Cell proliferation for both tumor cell lines were significantly reduced after the KD (*p=0.095, **p=0.087, t-test). **F.** Migration ability was assessed with wound-healing assays, with representative images captured at 0 h, 12 h, and 24 h. The wound gap area was quantified and normalized to 0 h. Significantly larger gap area for wound closure was observed in MDA-MB-231 after 24 hours of THAP2 KD (**p=0.05, t-test). For SKOV3, KD of ZIK1 resulted in significant bigger gap area after 12 hours (*p=8.21e-3, t-test). Scale bar is 200*μm*. **G.** Transwell migration assays were used to evaluate cell migration. Migrated cells were stained with crystal violet, and their total number was quantified. After KD, both cell lines showed lower number of migrated cells (*p=0.142 for MDA-MB-231, *p=0.05 for SKOV3, t-test). Scale bar is 200*μm*. **H.** Kaplan-Meier survival curves for THAP2 in TNBC patients, and ZIK1 in OV patients. In low-expression groups, both TFs showed higher survival probabilities (p=0.064 for TNBC, p=0.026 for OV). Error bars in **c**, **e**, **f**, and **g** denoted standard errors.

We then performed *in vitro* experiments to validate the new targets. Across various TNBC and OV cell lines, MDA-MB-231 and SKOV-3 cell lines were selected for further validations, as their expression profiles exhibited the highest consistence with the pseudobulk tumor-cell scRNA in the TNBC and OV tissues we used for predictions (**Fig. S7D**). For closest normal epithelial cells, MCF-10A and iOSE11 cell lines were used as normal mammary epithelial cells and normal ovary-derived secretory epithelial cells, respectively. The siRNA-mediated knockdown (KD) experiments of THAP2 in MDA-MB-231, and ZIK1 in SKOV3 were conducted, and over 65% KD efficiencies were observed in both cell lines (**Fig. 5C**). Next, bulk RNA-seq assays were used to measure gene expressions in both KD and control groups, and their cosine similarities with cancer and normal epithelial cell lines were calculated. For both MDA-MB-231 and SKOV-3, after 24 hours of KD of the targets, we observed significantly decreased cosine similarities with cancer cell lines, and increased cosine similarities with normal mammary or ovarian epithelial cells (**Fig. 5D**), which were consistent with the predicted cell states. In addition, significant reductions in cancer cell proliferation were achieved in both MDA-MB-231 and SKOV3 cell lines (**Fig. 5E**). Furthermore, wound healing assays and transwell migration assays were used to investigate the KD effects on cell migration. The cell migration abilities of the two cell lines were significantly reduced after the KD of our predicted TFs (**Fig. 5F, G**).

Finally, clinical relevance of THAP2 and ZIK1 were validated using Kaplan-Meier survival analysis (*91*) on public data. In TNBC patients with low THAP2 expression and OV patients with low ZIK1 levels, elevated survival probabilities were observed (**Fig. 5H**), validating the accuracy of PertFormer identified therapeutic targets.

## Discussion

In this study, we developed PertFormer, a foundational model with 3 billion parameters pretrained using extensive multiomic data. PertFormer enabled zero-shot predictions in regulatory and cellular tasks, specialized in perturbation studies. The novel architecture of PertFormer allows for pretraining on integrated multimodal of bulk and single-cell, total 9 types of multiomics datasets, facilitating the simultaneous extraction of comprehensive epigenetic and cellular heterogeneous features. PertFormer offers zero-shot simulations of a wide range of biological permutations. This capability makes PertFormer particularly valuable for downstream applications where experimental datasets are typically unavailable.

Diverging from transcriptome-focused models that typically rely solely on scRNA-seq data, PertFormer employed paired scATAC and scRNA as the pretraining data. By structuring each training sample around an individual gene, the pretraining of paired scATAC-seq and scRNA-seq data at gene level significantly expanded the effective sample size from the cell level (as in transcriptomic models) to the genic × cell level (as in our model).

We have systematically compared the state of the art models previously reported. Unlike those models focused directly on gene expression (*21, 23*), PertFormer performs *in silico* simulations by manipulating permutations on genomic and epigenomic regions. Through this process, not only can gene expression be accurately predicted, the capability inherent in PertFormer also enables the detection of causal regulatory relationships between epigenetic/regulatory interference and gene expression changes, providing interpretable insights of biological mechanisms. In contrast, transcriptomic models primarily capture correlative relationships among genes.

The ablation studies have demonstrated that increasing the model parameter size, the scale of pretraining datasets, and the input length can lead to improved performance of PertFormer (**Fig. S10**). With the rapid expansion of datasets generated from numerous biosamples across different species, as well as the swift advancements in computational resources, this foundational model can be further refined and adapted to more zero-shot tasks with enhanced performance.

## Acknowledgments

The authors thank Dr. Ken Chen for discussions and comments on model architecture and pretraining. The authors thank Dr. Xin Li for advice on statistical tests. The authors thank Junjie Wu for advice on training accelerations.

## Author contributions

Z.Y. and F.G. conceived and designed the study. X.L. and F.G. supervised the work. Z.Y. designed the model and developed the zero-shot methods. Z.Y. and F.G. contributed to implementation and application of the model. X.F. conducted *in vitro* validation of PertFormer-predicted treatment targets. Z.Y. and M.L. collected and integrated the datasets for pretraining. M.L., Z.Z. and B.L. performed analysis of other models used as comparisons in cell reprogramming. X.T. performed the survival analysis. Y.Y and L.T. performed analysis of other models used as comparisons in TF KI. Z.Y., X.F. and F.G. wrote the manuscript with the information from all authors. Z.Y., M.L., X.T., B.L., Z.Z., G.C., L.T., X.L. and F.G. edited and improved the manuscript.

## Data and materials availability

The codes of PertFormer are publicly available on GitHub: https://github.com/alibaba/damo-pertformer. The pretrained model parameters are publicly available on Hugging Face: https://huggingface.co/GenomicIntelligenceDamoAcademy/pertformer. We provide a web server (https://pertformer.ibreed.cn) for users to utilize PertFormer model.

## Materials and methods

### Pretraining datasets

#### Reference genomes

GRCh38 and mm10 were used for human and mouse genome assemblies, respectively. UCSC Lift Genome Annotations tool (*96*) was used to make conversions between various versions of reference genomes. For bulk data, GENCODE v29 and vM21 were used for human and mouse genome references according to the ENCODE4 pipeline (*97*). For single-cell data, GENCODE v32 and vM23 were used as the reference genome following the 10x Genomics 2.0.2 pipeline (*98, 99*).

#### Bulk epigenetic data for PertFormer-Elementary pretraining

Histone ChIP-seq data of H3K4me3, H3K4me1, H3K27ac, H3K27me3, H3K9ac, DNase-seq, and CTCF ChIP-seq data across 34 biosamples were collected from the ENCODE project for pretraining. ENCODE pipelines were applied to generate signal values from the bigWig files. Specifically, read-depth normalized signal was utilized for DNase-seq data, while signal p-value was used for all other epigenetic markers. ENCODE accession IDs of all these data were documented in **Data S1a**.

#### Single cell data for PertFormer-Regulatory pretraining

Single-cell multiomic data were gathered from public database and the accessions were documented in **Data S1b**. Filtered feature barcode matrix, cell barcodes, gene lists, and consensus ATAC peak coordinates were downloaded, and Scanpy 1.9.8 (*100*) was used to form the raw scRNA-seq and scATAC-seq matrices. The scATAC-seq signals were generated through the following steps: (1) downloading the ATAC per fragment information file (.tsv.gz file) and splitting it by cell barcode to obtain the fragment information (.bed file) for individual cells; (2) converting the bed file into bam format for each cell using bedtools 2.26.0 (*101*); (3) using bamCoverage from deepTools 3.5.4 (*102*) with counts per million (CPM) normalization to obtain the scATAC signal for each single cell.

#### bulkMultiomeCorpus-55b

For each biosample, 50,000 × 32,000-bp non-overlapped bins were randomly sampled from the GRCh38 assembly. DNA sequences information, along with the corresponding 7 types of epigenetic signals, were collected from each bin. Specifically, epigenetic signals were collected at 1-bp resolution. In total, the 34 biosamples comprised a bulk multiomic dataset for pretraining, with a cumulative length exceeding 55 billion bp, which was approximately 17 times the length of the human genome.

#### scMultiomeCorpus-1.5b

Scanpy was used for the quality control (QC) of the paired scRNA-seq and scATAC-seq data. Briefly, genes that were expressed in fewer than 3 cells and cells that expressed fewer than 200 genes were removed. Cells that expressed too many genes or had high mitochondrial genes counts were also removed. For scATAC dataset, cells with a unique peak number less than 0.5% of all consensus peaks were excluded. The intersection of remaining cells respectively in scRNA and scATAC data were eventually used. Gene expression values were adjusted using total counts normalization, followed by *log*(1 + *x*) transformation. Highly variable genes were identified by use of Scanpy. In each biosample, genes that were highly variable and expressed in more than 10% of all single cells were selected for pretraining. In addition, genes with fewer than 4 consensus chromatin-accessible peaks flanking 300,000-bp of the TSS, or with unknown nucleotides in any of the consensus peaks in the same 300,000-bp range, were excluded. The remaining genes for each biosample were listed in **Data S1c**, which covered over 15k genes. In our dataset, each sample represent a gene in a single cell, which contained multiomic information (DNA sequences and scATAC signals) of all consensus peaks flanking 300,000-bp of its TSS, along with the gene expression value. For each consensus peak, the 1,024-bp region surrounding the peak center was used. Overall, the number of used genes, when multiplied by the number of used cells, led to more than 1.5 billion samples, forming our scMultiomeCorpus-1.5b pretraining dataset.

### PertFormer architecture

#### Data pre-processing

K-mer tokenization (*103*) strategy was used to convert DNA sequences into word tokens. Here *K* = 6, leading to 4^6^ = 4096 unique DNA tokens. For epigenetic signals, we utilized a categorical discretization method to tokenize signal values by use of a set of 36 distinct word tokens (*104*) (**Data S1d**). Besides, we applied two special tokens, [MASK] and [CLS]. The [MASK] token was used to represent the masked DNA or epigenetic tokens during the pretraining of bulk data. The final hidden state of the [CLS] token can be used as the aggregated representation of a multiomic sequence (*105*). We added the [MASK] token to both DNA vocabulary and epigenetic vocabulary. [CLS] token was added to the DNA vocabulary. Consequently, tokens in the DNA vocabulary (*Token*_*DNA*_) can be encoded using integers ranging from 0 to 4097, while tokens for each kind of epigenetic signals (*Token*_*epigenetic*_) can be represented by integers from 0 to 36.

#### PertFormer-Elementary, Multiomics embeddings layer

8 trainable embeddings (*106*) were assigned respectively for DNA sequences and 7 types of epigenetic signals, each projecting tokens into vectors with a dimensional size of 2048 (*d*_*model*_). The multiomic embeddings were obtained by direct summation of all these 8 embeddings. Specifically, for epigenetic signals not included in the input, their corresponding embeddings will be set to zero. PertFormer-Elementary accepts input lengths of 128, 1,024, 2,048, 4,096, 8,192, or 16,384 bp. The multiomic embeddings were divided into 128-bp bins. At the beginning of each 128-bp bin, we additionally assigned a [CLS] token projected by the DNA embedding (*E*_[*CLS*]_). Integers from 0 to 128 were employed to represent the 1+128 positional tokens in each bin (*Token*_*position*1_), which were projected into 2048 dimensions using Position-1 embedding layer (a trainable embedding). The final embeddings fed into the following transformer blocks were the direct summation of the multiomic embeddings and the Position-1 embeddings. The Multiomic embedding layer therefore contains *d*_*models*_ × (*Token*_*DNA*_ + *Token*_*epigenetic*_ × 7 + *Token*_*position*1_) = 9, 187, 328 parameters.

#### PertFormer-Elementary, Transformer-1 Block

The Transformer-1 Block was comprised of 20 Transformer encoder layers (*107*). Each layer was structured with a multi-head attention (MHA) operation, followed subsequently by a feed-forward unit. Given the embedding sequence denoted as *E*, the *MHA* operation can be executed as

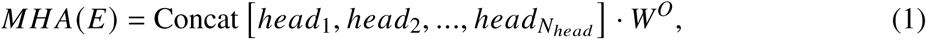

where

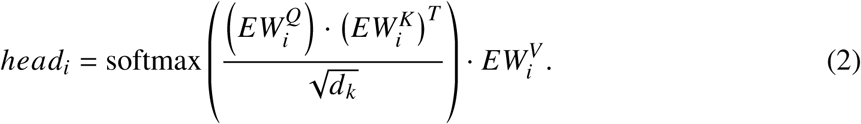

Here, 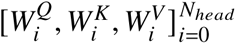 were trainable linear projections that reduce the dimension (from 2048 to 64), while the trainable linear projection *W*^*O*^ recovered the concatenated multiple heads back to the shape of input embeddings. In the MHA operation, we set the dimension of K, Q, V to 64 (*d*_*head*_) and the number of head to 32 (*N*_*head*_), and used hidden neurons numbering *d*_*model*_ ×4 = 8192 (*d*_*hidden*_) in the feed-forward units. Hence, the total parameters *P*_*total*_ in Transformer-1 Block in PertFormer-Elementary can be calculated as

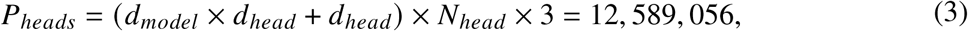

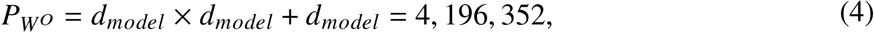

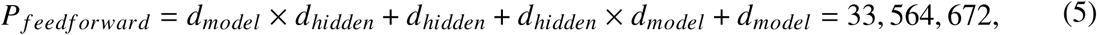

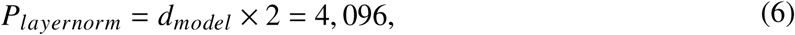

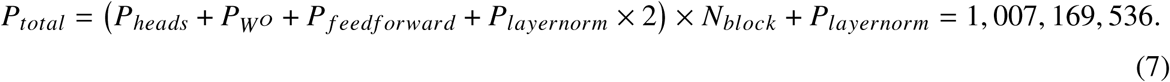

Transformer-1 Block was deployed to process each bin of multiomic embeddings. In Transformer-1 output of each bin, the final hidden state of its [CLS] token (*R*_[*CLS*]_) was used as the representation of this bin, and was fed into the next stage.

#### PertFormer-Elementary, Transformer-2 Block

The *R*_[*CLS*]_ vectors from all bins were concatenated together. Similar to Position-1 embedding, we used consecutive integers starting from 0 to denote the positional token of each bin and applied Position-2 embedding layer to project them into 2048 dimensions. Position-2 embedding has *d*_*model*_ ×128 = 262, 144 parameters. The direct summation of concatenated *R*_[*CLS*]_ vectors and Position-2 embeddings were fed into Transformer-2 Block. Transformer-2 Block was of the same structure as Transformer-1 Block (also with 1,007,169,536 parameters). The output of Transformer-2 Block extracts mid-range information from all bins, forming the overall characteristics of the entire genomic elements.

#### PertFormer-Regulatory, processing each peak

PertFormer-Regulatory module targets each gene individually, and its parameters were shared across all genes. For each gene, the regulatory element candidates (RECs) were consensus scATAC peaks within a 300,000-bp range flanking the TSS. The lengths of scATAC peaks were resized to 1024 bp centered at the peak summit. The input information for each REC comprised DNA sequences and scATAC-seq signals, which were processed by PertFormer-Elementary. Next, the output vector of Transformer-2 for each REC went through an attention pooling process. Attention pooling was defined as follows:

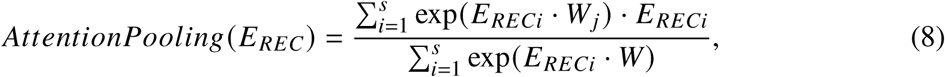

where *E*_*RECi*_ represented the *i*^*th*^ vector in the REC embedding, *s* was the pooling size and was set to 8, and *w* ∈ *R*^2048×*s*^ was a matrix of learned weights. All REC vectors were concatenated to form a long-range multiomic vector with a size of (*P*, 2048), where *P* was the number of input RECs. The attention pooling module therefore has *d*_*model*_ × *d*_*model*_ = 4, 194, 304 parameters.

#### PertFormer-Regulatory, Transformer Block

We defined a series of numbers to represent the distance of each REC from the TSS. The REC closest to the TSS was assigned with a value of 0. Increased continuous odd or even number was used to respectively represent the REC that was on the upstream or downstream of TSS, ascending by the distance to TSS. These numbers were then projected through a TSS-distance embedding layer and added to the long-range multiomic vector of the RECs. For CREforemr, the maximum number of RECs is set to 150. Therefore, the TSS-distance embedding layer has *d*_*model*_ × 150 × 2 = 614, 400 parameters (since odd and even numbers were used, the input tokens were doubled). The Transformer Block in PertFormer-Regulatory was also of the same structure as the Transformer-1 Block (also with 1,007,169,536 parameters), and the output of this block forms the multiomic learning representation of all regulatory elements of each specific gene.

#### PertFormer-Regulatory, sc/bulk gene expression prediction

Since the numbers of RECs varied among genes, zero-padding was used to make sure the outputs were of the same shape. The padded outputs were then flattened and fed into a 2-layer fully connected feed-forward network, with one hidden layer of 32 hidden neurons with ReLU activation function (*108*), followed by the output layer with one neuron to predict the final gene expression. Specifically, all outputs were zero-padded to a fixed length of 150 RECs. Therefore, the feed-forward network has *d*_*model*_ × 150 × 32 + 32 + 32 × 1 + 1 = 9, 830, 465 parameters.

Overall, the total parameters in PertFormer were 9, 187, 328 + 1, 007, 169, 536 × 3 + 262, 144 + 614, 400 + 4, 194, 304 + 9, 830, 465 = 3, 045, 597, 249, which is over 3 billion.

### PertFormer pretraining

#### PertFormer-Elementary pretraining, mask-reconstruction method

Firstly, the PertFormer-Elementary module was pretrained on bulkMultiomeCorpus-55b. For each training batch, input multiomics with lengths of 128, 1,024, 2,048, 4,096, 8,192, or 16,384 bp were randomly sampled from the 32,000-bp training data bins. From these samples, 1-7 types of input epigenetic signals were randomly selected, along with DNA sequences, forming the input multiomics (referred to as the Selected group). The rest of the epigenetic signals were the Preserved group. Within each 128-bp bin, the mask reconstruction method was performed through the Transformer-1 Block. Specifically, 3 distinct DNA sequence segments (each consisting 6 consecutive tokens), and a segment with 64 continuous tokens in one of the epigenetic tracks from the Selected group, were masked. All those segments and tracks were selected at random. To perform the mask, the corresponding tokens were replaced with [MASK] tokens. To predict these masked tokens, 8 classification modules were deployed respectively for the DNA sequences and the 7 kinds of epigenetic signals. For each masked token, its output vector from Transformer-1 Block was fed into the corresponding classification module. Specifically, each classification module was a 2-layer fully connected feed-forward network, with 2048 input neurons, one hidden layer of 32 ReLU-activated hidden neurons, and 4096 output neurons for DNA sequence classification or 36 output neurons for epigenetic signal classification. Cross-entropy loss was employed.

#### PertFormer-Elementary pretraining, prediction of Preserved multiomics method

The output vectors from the Transformer-2 Block were utilized to predict the values of the multiomic signals in the Preserved group. For each 128-bp bin, the average signal value of each epigenetic information in the Preserved group was calculated, discretized into word tokens, and used as ground truth. Seven classification modules, each identical in architecture to those in the mask-reconstruction method, were constructed after Transformer-2 Block to predict the epigenetic signals in the Preserved Group. Cross-entropy loss was employed.

#### PertFormer-Elementary pretraining hyper-parameters

During PertFormer-Elementary pre-training, the total loss was the sum of the losses from the two pretraining methods. Throughout this study, unless otherwise specified, the ADAM optimizer was used to update the neural network weights, distributed data parallel strategies were applied to do the training on 32 NVIDIA A100 GPUs, and the ‘bfloat16’ data type in the torch.cuda.amp function was used to accelerate the training. In the initial training stage, the learning rate was set to 1e-5, and was gradually reduced to 5e-7 during the training. Early stopping was adopted when the loss value converged (no further reduction was observed for five consecutive epochs).

#### PertFormer-Regulatory pretraining in scMultiomeCorpus-1.5b

All parameters in the CREformer-Elementary module were frozen during this pretraining. In scMultiomeCorpus-1.5b, for each gene in each single cell, PertFormer was pretrained to predict the gene expression level based on DNA + scATAC information of all consensus peaks flanking 300,000-bp of its TSS. The mean squared error (MSE) loss function was used for gene expression regression.

#### PertFormer-Regulatory pretraining hyper-parameters

In the initial training stage, the learning rate was set to 1e-6, and was gradually reduced to 1e-7 during the training. We retained 10% of single cells in scMultiomeCorpus-1.5b for validation, and another 10% of single cells for test. Early stop was made when the PCC of single-cell gene expression predictions converged in the validation data.

### Zero-shot predictions of master regulators and GRNs

#### Datasets

The 10× scMultiome datasets of PBMC (pretrained cell prediction) and CRC tumor (*109*) (unseen cell prediction, **Data S2c**) were used. The data pre-processing protocols were similar to the scMultiomeCorpus-1.5b. Specifically, since PertFormer predictions were solely based on scATAC and DNA sequences, scRNA data were assumed to be unavailable. Hence, for model input, only scATAC QC steps were performed. Notably, scRNA QC steps were performed for scRNA control groups of cell clustering and master regulator predictions, but without cell filtering in order to keep the same remained cells as scATAC for evaluation.

#### Master-TF correlations

We defined the Attention Score on the *j*^*th*^ peak as

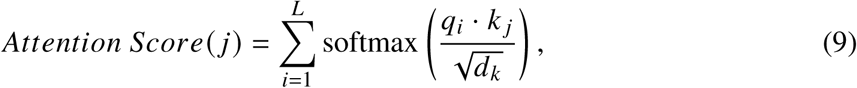

where *q*_*i*_ and *k* _*j*_ represent the *i*^*th*^ and the *j* ^*th*^ vectors from the Queries and Keys matrices, respectively. *L* denotes the number of RECs. Here we used the mean Attention Score across all heads and layers in PertFormer-Regulatory Transformer Block.

We defined B cells of PBMCs (included in scMultiomeCorpus-1.5b) as the pretrained cells group and tumor cells of CRC (not included in scMultiomeCorpus-1.5b) as the Excluded-cells group. Both groups incorporated genome-wide genes. For each of the two groups, Spearman correlation coefficients (SCC) between master regulator ChIP-seq signals and Attention Scores at RECs were calculated. For example, for each gene in the pretrained cells group, the mean Attention Score across all B cells were generated, and the SCC between this averaged Attention Score and each of the three B cell master regulators’ ChIP-seq signals (PAX5, EBF1, and POU2F2, ChIP-seq signals collected in GM12878 cell line from ENCODE) were computed respectively. Averaged SCC among the three TFs were reported. For control group, SCC were calculated between master regulator ChIP-seq signals and the aggregated scATAC signals in the B-cell cluster at RECs. Similarly, for genes in the unseen cells group, the correlations were measured between master regulator and the averaged Attention Score or aggregated scATAC signals in CRC tumor-cell cluster.

#### Single-cell clustering based on gene expression or Attention Feature

We defined the Attention Feature as follows:

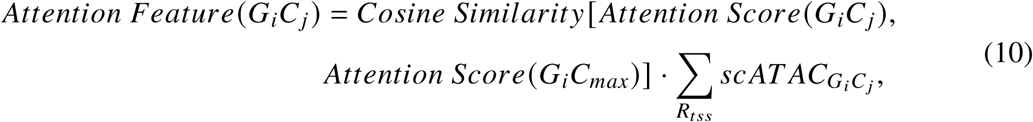

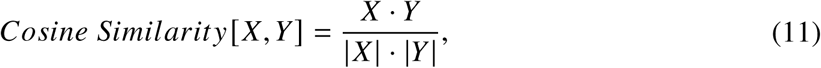

where *G*_*i*_*C*_*j*_ denotes the *i*^*th*^ gene in cell *j* ^*th*^, and *G*_*i*_*C*_*max*_ denotes the *i*^*th*^ gene in the cell with the maximum value of total scATAC signals in all RECs of this gene. 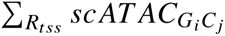 denotes the total scATAC-seq signals within the consensus peaks flanking 20,000-bp of the TSS of the *i*^*th*^ gene in *j* ^*th*^ cell. Attention Feature extracted multiomic features of each gene in every cell.

The following clustering processes were conducted by Scanpy, using either gene expression or Attention Feature as input data. Briefly, the effects of total counts per cell and the ratio of expressed mitochondrial genes were regressed out, and the data were normalized and truncated with a maximum value of 10. Principal component analysis (*110*) (PCA) was performed using the “arpack” solver. The nearest neighbors distance matrix was computed to generate neighborhood graph. Cell clusters were identified using Leiden methods (*111*). The clustering results were visualized on UMAP.

#### Single-cell cell type annotations

Marker genes (*109*) were utilized to annotate the cell type (**Data S2d**). For each marker gene, Scanpy was employed to generate and visualize the fraction of cells, along with the mean expression or the mean Attention Feature in each cell cluster.

#### Predictions of cell type-specific master regulators

After cell type annotations, Scanpy (with ‘logreg’ as the method (*112*)) was used to identify TFs with high and differential Attention Features, defined as master regulators. We selected 6 highly-ranked regulators for each cell type, and plotted the fraction of cells and mean Attention Feature. For control groups, scRNA or scATAC profiles were used. For scRNA, the Attention Feature of each gene in each cell was replaced by its gene expression, and therefore differentially expressed TFs were identified. For scATAC control, the Attention Scores for calculations of Attention Features (**Formula** 10) were replaced by the scATAC signals.

**GRN inferences.**

For each cell type, GRNs were constructed between master regulators and their targeting genes. For a specific gene in a cell type, PertFormer-Regulatory was used to generate the mean Attention Score across all cells of that cell type, and the list of top 15% RECs ranked by Attention Score (or at least 2 RECs) were prioritized for further motif analysis. RECs that covered the TSS were excluded from the list. FIMO (*113*) was utilized to scan the motifs of master regulators on the list of top RECs with default parameters. The motif enrichment was defined as

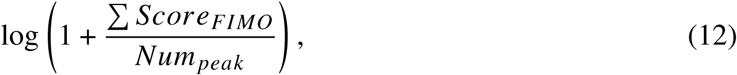

where *Score*_*FIMO*_ corresponds to the 7^*th*^ column in FIMO’s result table. A higher motif enrichment indicates a greater likelihood that this master regulator is regulating the gene. For comparison, GRN predictions were performed on the same group of genes collected from the SCENIC+ study. To construct the GRNs of B cells, we calculated the motif enrichment of 3 master regulators on those genes, and annotated the GRN manually. The consistency of GRN inference between PertFormer and SCENIC+ was assessed by measuring the proportion of genes that were categorized into the same group by both models across the seven regulatory combinations. For each TF, every gene linked to this TF in the predicted GRN maps formed a TF-to-gene relationship. We also assessed the proportion of these links in SCENIC+ that also appeared in PertFormer’s prediction. In the CRC case, GRN inference was conducted on the top 100 DEGs identified by Scanpy in tumor cells, based on PertFormer-predicted gene expressions. For both tasks, genes without significant motif enrichment for all 3 TFs were excluded from the analysis.

### Zero-shot predictions of perturbations on transcriptional regulation

#### Datasets

In the K562 CRISPRi-FlowFISH study (*114*), CRISPRi-validated significant targets with regulatory distances less than 150,000 bp were collected (**Data S3a, b**). Specifically, both up-regulated (RG-pairs) and down-regulated (EG-pairs) expressions were observed across different significant CRISPRi targets. For EG-pair predictions, validated EG-pairs and the non-significant pairs were used. For perturbation simulations on EG-pairs and RG-pairs, all CRISPRi-validated significant pairs were used. For prediction of EG-pairs in mouse tissues, all *in vivo* validated true and false EG-pairs with regulatory distances less than 150,000 bp were collected from the VISTA mouse enhancer database (*115*) (**Data S3c**). For TF perturbations, KO datasets were collected from the ENCODE project (**Data S4**), and KI datasets were collected from previous study (*116*).

#### Predictions of EG-pairs in K562 CRISPRi-FlowFISH dataset

Bulk ATAC-seq profiles from K562 cell line (**Data S3d**), along with the DNA sequences, were utilized as input information. For each gene in the EG-pairs, any bulk ATAC peak flanking 300,000-bp of the TSS was considered as its REC, and the Attention Scores were calculated. The RECs were then individually ranked for each gene by their Attention Scores, with the final ranking for each REC defined as its rank divided by the total number of RECs for that gene. Violin plots were used to present the ranking in both positive and negative EG-pairs. As control, rankings based on ATAC-seq signal were generated. Based on the final ranking, we used area under precision-recall curve (auPRC) to evaluate the classification performance across different regulatory distances. Besides ATAC signals, we used H3K27ac (*117*), linear regression (*118*), Enformer (*119*), SCARlink (*120*), and GET (*121*) as comparison methods, and related implementation details were provided in **Supplementary Text**.

Notably, throughout this study, unless otherwise specified, the methods for using bulk data as open chromatin input were similar to this section.

#### Predictions of EG-pairs in mouse VISTA database

The coordinates were converted from mm9 to mm10 assembly using UCSC Lift Genome Annotations tool. Bulk ATAC-seq of mouse tissues were obtained from the ENCODE website (**Data S3d**). Other experimental settings were similar to the K562 CRISPRi-FlowFISH dataset.

#### Predictions of perturbations on EG-pairs and RG-pairs

For all CRISPRi-validated significant pairs in K562 FlowFISH dataset, to perform the *in silico* perturbations on enhancers or repressors, original (Ref) and removed (KO) ATAC-seq signals were respectively applied to the RECs co-localized with the enhancer or repressor region. The original ATAC signals were retained for the remaining RECs in both KO and Ref conditions. Gene expression levels were then predicted for both KO and Ref, and their differences were calculated. Accuracy and macro-F1 were measured after *in silico* KO by PertFormer, in which perturbation targets with decreased expression levels of targeted genes were predicted as EG-pairs, and vice versa for RG-pairs. Besides, PCC_delta was measured as the PCC (*122*) between predicted and experimental gene expression changes. In **Fig. 3E**, fraction change refers to the percentile of the change in gene expressions:

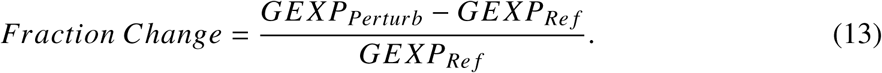

#### Predictions of TF KO perturbation

The experimental perturbation operation of this dataset was CRISPRi, shRNA (*123*) or siRNA (*124*) in K562 cell line. Bulk ATAC-seq profile was used as the open chromatin information. For each of the perturbed TFs in the dataset, FIMO program was employed to scan the motif in all ATAC-seq peaks. Peaks with motif p-value less than 1e-5 were selected to perform the perturbation. The ATAC-seq signals in all of these selected peaks were set to zero to simulate the TF deletions. Original (Ref) and perturbed (KO) ATAC-seq signals were employed as the open chromatin information for PertFormer to generate the predicted gene expression before and after the *in silico* perturbations, together with the gene expression difference. Distributions of predicted gene expression changes were evaluated on DEGs identified by previous work (*125*). In these DEGs, macro-F1 was evaluated by comparing the concordance of predicted and experimental gene expression changing directions, and PCC_delta was measured between the predicted and experimental gene expression changes.

#### Predictions of TF KI perturbation

This dataset was generated from the perturb-seq gene activation experiments (*116*) in K562 single cells, and only TF KIs were used. We employed bulk ATAC profile as the chromatin accessibility information. For each of the perturbed TFs, on genome-wide scope, FIMO with default parameters was used to scan for its motif. Regions of 1024 bp flanking the center of each FIMO-detected TF motif were defined as motif loci. For each gene, motif loci with the highest FIMO score (could be multiple if they shared the highest FIMO score) flanking the 300,000-bp of its TSS were selected as KI loci, and artificial chromatin-accessible peaks were added to the selected loci to perform the *in silico* KI. Specifically, the signal information of the artificial peaks was directly copied from the highest ATAC peak among all peaks flanking the 300,000-bp of the TSS of that gene. Next, PertFormer was used to predict gene expression levels before (Ref) and after (KI) the perturbation. Notably, co-overexpressions of multiple TFs were also included in the KI datasets. For *in silico* overexpression of multiple TFs, KI loci were calculated independently for each TF, while the ATAC signals were added to all the KI loci for all TFs simultaneously. For evaluation, DEGs were identified using Scanpy (with ‘t-test’ as the methods) between the perturbed and wild-type K562 cell clusters, and macro-F1 was used to assess the predicted gene expression changing directions of the top 200 up- and 200 down-regulated DEGs. PCC_delta was computed on all genes. Here, the experimental delta values were defined as the changes of averaged gene expressions in the KI and Ref cell clusters:

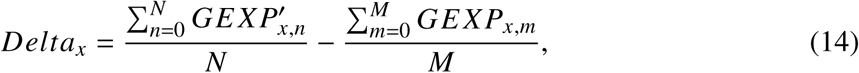

where *GEXP* and *GEXP*^′^ respectively refers to the expression profiles of the wild-type and perturbed cell clusters, *x* refers to the specific gene, *N* and *M* respectively refers to the total number of cells in the wild-type and perturbed cell clusters.

SCENIC+ (*125*), scGPT (*126*), and scFoundation (*127*) were used as comparisons in the TF perturbation tasks. We also performed control experiments, in which KO and KI were performed at random loci without considering the TF motifs. Related details were provided as **Supplementary Text**.

### Zero-shot Predictions of cell state transitions

#### Dataset

The epigenetic and transcriptomic data of hESC, fibroblast, cardiomyocytes, and B cells were collected from the ENCODE project (**Data S6a** and **Data S6b**). Briefly, for bulk RNA-seq data, the adjusted (by log(1 + *x*)) Transcripts Per Kilobase Million (TPM) profiles were used to represent gene expression values. For bulk chromatin accessibility profiles, bulk ATAC-seq or DNase-seq profiles were used, depending on the availability of experimental data. If both open chromatin markers were available, ATAC-seq was used. For single-cell data, the mouse embryonic brain E18 scMultiome dataset were collected from the 10x Genomics website. DEGs from hESC to somatic cells were used for evaluations on bulk data, which were screened out by the following steps: (1) genes were ranked by log_2_ *FC* between the two stages in descending order, and genes with log_2_ *FC* > |1| were retrained; (2) genes from (1) were ranked by absolute difference of gene expression in descending order; (3) intersection of top 50% genes in (1) and top 50% genes in (2) were used. For mouse reprogramming simulations, all genes with log_2_ *FC* > |1| were used as DEGs.

#### Predictions OSKM KI to bulk somatic cells

For each of the 3 somatic cells, co-overexpression of SOX2, OCT4, KLF4, and c-MYC were simulated through PertFormer *in silico* KI. The KI methods were described in the previous “Predictions of TF KI perturbation” section. Accuracy and macro-F1 were evaluated on the predicted gene expression changing directions of the identified DEGs. Here, according to the reprogramming effects, decreased gene expressions were considered ground truth for up-regulated DEGs from hESC to somatic cells, and vice versa. Cosine similarity was used to compare the gene expression distributions of DEGs between cell states. For PertFormer predicted gene expression of somatic cell or reprogrammed somatic cell, its cosine similarities with experimental gene expression of somatic and hESC states were evaluated, respectively. Besides, in RNA-seq profiles, cosine similarities between somatic and hESC states were also measured. Geneformer and scFoundation were used as methods for comparison, and the implementation details can be found in **Supplementary Text**. Notably, since Geneformer outputs the rank of gene expression which does not imply increased or decreased gene expressions, it is not able to predict the changing directions of DEGs (**Fig. 5C**).

#### Simulations of different OSKM TF KI combinations

The DEGs identified between fibroblast and hESC were used as input features for cell clustering. The profiles of experimental gene expression differences were used in PCA analysis. Fibroblast was set as the reference cell to be compared, and the profile of gene expression differences **D** in hESC or fibroblast can be represented as

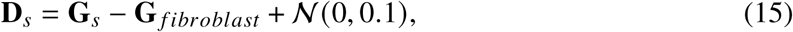

where *s* denotes either hESC or fibroblast, **G**_*s*_ − **G** _*fibroblast*_ denotes the expression differences of all DEGs, N (0, 0.1) is a noise of normal distribution, with zero mean value and 0.1 variance. Since we only have 2 cells with experimental data, which was not enough for PCA analysis, we expanded the number of samples by repeating the **Equation.**15 for 1000 times for each of the 2 cells, leading to 2000 samples in total. Scanpy was used with processes similar to previous GRN inference workflow to perform the PCA analysis.

ATAC-seq in fibroblast was used as the reference (Ref) chromatin accessibility profiles (**Fig. 5F**). The 4 OSKM TFs formulated 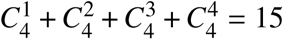 combinations, while ATAC KI process was performed independently for each combination. The UMAP visualization of fibroblast, hESC, and the 15 reprogrammed cells were formed by use of “sc.tl.ingest” function. Specifically, we used **D**_*s*_ = **G**_*s*_ − **G** _*fibroblast*_ (noises removed) as input features of fibroblast and hESC, while gene expression differences between KI and Ref were used as input features of the reprogrammed cells.

#### *In silico* reprogramming using single-cell data

Highly-variable genes (HVGs) from scRNA profiles of mouse embryonic midbrain E18 were determined by Scanpy, and were used for analysis. Single-cell clustering and cell type annotation were performed based on the Attention Feature, following the GRN inference workflow. For each HVG in every NPC cell (Ref), *in silico* KI of OSKM was performed, and Attention Features were re-calculated. The “sc.tl.ingest” function was used to form the UMAP plot of all original and *in silico* reprogrammed cells. This function also assigned cell type labels for the reprogrammed cells, which were used to calculate the percentile of cells that reached the ESC state. The directions of cell state transitions were marked with yellow arrows. Geneformer and scFoundation were used for comparison and related details were in **Supplementary Text**.

### Zero-shot predictions of novel targets

#### Single-cell clustering and cell type annotations

Following PertFormer’s zero-shot GRN inference workflow described in the “Zero-shot predictions of master regulators and GRNs” section in **Materials and methods**, single-cell clustering was performed based on the Attention Feature for each of the 7 types of tumor tissues collected from previous work (*109*) (**Data S2c**). Closest normal cells (CNCs) of these 7 tumor types were defined by previous studies (*109*): luminal mature cells for BRCA of the non-basal subtypes, luminal progenitor cells for TNBC, distal stem cells for CRC, secretory endometrial epithelial cells for OV and UCEC, ductal-like-2 cells for PDAC, and melanocytes for SKCM. Cell types were annotated by Attention Feature of marker genes (**Data S2d**). In addition, cell clustering and cell type annotations were also performed in each of the 7 tumor tissues using scRNA data. In comparison between PertFormer’s zero-shot predictions and scRNA’s results, cell type annotations yielded 80.33% consistency, on average across the 7 tumor tissues.

#### Predictions of tumor-specific master regulators

For each of the 7 types of tumor tissues, with input of the corresponding scATAC profiles, along with the DNA sequences, master regulators were identified following PertFormer zero-shot GRN inference workflow. The tumor-promoting and tumor-suppressive effects of the predicted regulators were collected from previous studies, which can be found in **Supplementary Text**.

#### *In silico* perturbations of predicted master regulators

For perturbation simulations in the 5 types of tumors in **Fig. S6**, top 1 tumor-promoting and top 1 tumor-suppressive TFs were used. For **Fig. S7**, all TFs that ranked before the novel targets were simulated. Specifically, since the TET3 motif is not available, and the regulatory effects of USF3 in OV is so far not conclusive, these two TFs were excluded from the simulations. Pseudobulk expressions were used to determine the use of KO or KI perturbations: for each TF, if its mean expression of tumor cells is higher than normal cells, KO perturbation will be adopted, and vice versa for KI perturbation. For unperturbed cells, single-cell clustering and cell type annotations were performed based on Attention Feature predicted from the reference scATAC-seq profile, following the GRN inference workflow. The KO or KI perturbations were performed on all tumor cells, following the protocols described in the “Zero-shot predictions of epigenetic perturbations” in **Materials and methods**, and their Attention Features were re-calculated after the perturbations. We used “sc.tl.ingest” function to form the UMAP plot of all original and *in silico* TF-KO cells. The directions of cell state transitions were marked by yellow arrows.

#### Identification of novel treatment targets

Following the rank of master regulators predicted by PertFormer, we searched for the first TF that has never been reported by other studies in the corresponding tumor context. Therefore, THAP2 was identified in TNBC and ZIK1 was identified in OV. The methods for perturbation simulations were similar to the previous descriptions.

#### Cell line selections

To select the closest cancer cell lines for *in vitro* experiments, bulk RNA-seq profiles of 3 TNBC cell lines (MDA-MB-231, HCC19377, BT-549), and 4 OV cell lines (SKOV3, OVCAR-3, ES-2, A2780), were collected from public database. Their accession IDs were specified in **Data S2e**. For bulk profiles, TPM of each gene was used. In the scRNA profiles of the selected tumor tissues, we calculated the pseudobulk expression of each gene in the tumor cells. PCC and cosine similarity were measured on genome-wide genes, between each cancer cell line and the tumor single cells.

For closest normal epithelial cells, MCF-10A cell line was selected as the normal mammary epithelial cells, while iOSE11 was used as the normal ovary surface epithelial cells. Their accession IDs were provided in **Data S2e**. TPM values were used.

#### Cell culture

Cancer cell lines, including MDA-MB-231 and SKOV3, were cultured under specific conditions: MDA-MB-231 cells in DMEM supplemented with 10% FBS, and SKOV3 cells in McCoy’s 5A medium (16600082, Gibco) with 10% FBS. All cancer cell cultures also included 1% penicillin-streptomycin. Cell lines were obtained from ATCC and maintained at 37^◦^*C* in a humidified atmosphere with 5% *CO*_2_.

#### Quantitative real-time PCR (RT-qPCR)

Total RNA was extracted using the FastPure Cell or Tissue Total RNA Isolation Kit V2 (RC112-01, Vazyme) and reverse transcribed into cDNA using the ABScript III RT Master Mix (RK20429, ABclonal). Quantitative RT-PCR was performed on 5*ng* of cDNA with the 2X Universal SYBR Green Fast qPCR Mix (RK21203, AB-clonal) in a 10*μL* reaction volume. The primers used were as follows: GAPDH (forward: 5’-GGAGCGAGATCCCTCCAAAAT-3’; reverse: 5’-GGCTGTTGTCATACTTCTCATGG-3’), ZIK1 (forward: 5’-AGGACATCGCCATTTACTTCTCA-3’; reverse: 5’-GCATCACTTCAAGGTACAG-GAG-3’), and THAP2 (forward: 5’-AGAATGGGTTCGCCTGGTTAG-3’; reverse: 5’-TGTTAG-GTCAAAACAGGAGGCT-3’). Cycling conditions were performed according to the manufacturer’s instructions for the SYBR Green Fast qPCR Mix, and amplification signals were detected using the Qtower 384G system (Jena). Gene expression levels were normalized to GAPDH Ct values, and the relative expression for each group was calculated using the 2^−ΔΔ*Ct*^ method relative to the control group.

#### Gene silencing

Small interfering RNAs (siRNAs) were purchased from Tsingke Biotech, with three siRNAs synthesized for each target gene and mixed to induce gene silencing. The sequences of the siRNAs were as follows: for siTHAP2 (sense-1, 5’-CCCUAUAGAAAUAUGUGUA-3’; antisense-1, 5’-UACACAUAUUUCUAUAGGG-3’; sense-2, 5’-GAACGCAGAGCAACUCGAA-3’; antisense-2, 5’-UUCGAGUUGCUCUGCGUUC-3’; sense-3, 5’-GACCUAGCUUAGAUAAA-UG-3’; antisense-3, 5’-CAUUUAUCUAAGCUAGGUC-3’), and for siZIK1 (sense-1, 5’-GCAUAC-GUGUGACCCAGUU-3’; antisense-1, 5’ AACUGGGUCACACGUAUGC-3’; sense-2, 5’GAAAC-CCGGCACAAUUACU-3’; antisense-2, 5’-AGUAAUUGUGCCGGG-UUUC-3’; sense-3, 5’-CA-ACACCGAAGAAUUCACA-3’; antisense-3, 5’-UGUGAAUUCUUCGGUGUUG-3’); A non-targeting control siRNA with no homology to any human genome sequence was also included as negative control. Transfection was performed using jetPRIME reagent (101000046, Polyplus) following the manufacturer’s protocol. siRNAs were diluted in jetPRIME buffer, with 0.3*μL* of jetPRIME reagent added to 10*μL* of buffer, or 1.5*μL* added to a 50*μL* buffer volume. cells were refreshed with antibiotic-free medium 24 hours prior to transfection.

#### Bulk RNA Sequencing

MDA-MB-231 and SKOV3 cells were cultured in 6-well plates and transfected with either non-targeting control siRNA or gene-specific siRNAs targeting THAP2 and ZIK1, respectively, using the jetPRIME transfection reagent following the manufacturer’s recommended protocol. Cells were collected at 24 hours post-transfection. Following collection, cells were washed twice with PBS and lysed directly in TRIzol Reagent (Thermo Fisher Scientific, #15596018). Lysates were stored at -80°C until RNA extraction. Total RNA was isolated from each sample, and polyadenylated mRNA was selectively enriched for library preparation. The resulting cDNA libraries were sequenced on an Illumina NovaSeq 6000 platform according to standard sequencing protocols.

#### Bulk RNA-seq data processing

Bulk RNA-seq fastq files were processed by Salmon v1.10.3. The GRCh38 reference genome and the index file were collected from NCBI database (**Supplementary Text**). Pytximport v0.12.0 was used to summarize transcripts to gene-level counts. TPM values were calculated and used for analysis. For calculations of cosine similarities, DEGs with *log*_2_*FC* > |1.5| were used.

#### Cell proliferation and migration assay

Cell proliferation was assessed using the Cell Counting Kit-8 (CCK-8, CK001, Lablead). A total of 5,000 cells were seeded into each well of a 96-well plate. After 48 hours of siRNA transfection, 10*μL* of CCK-8 solution was added to each well and incubated at 37^◦^*C* for 2 hours. Absorbance at 450*nm* was measured using the Cytation 5 Cell Imaging Multi-Mode Reader (Agilent). A standard curve correlating cell number to 450*nm* absorbance was established for each cell line and subsequently used to calculate cell counts. Cell counts were normalized to the control group, which was transfected with negative control siRNA. For the wound healing assay, 5 × 10^4^ cells were seeded into a 24-well plate. After 24 hours of siRNA transfection, an artificial wound was created using a 200*μL* pipette tip. Wound closure was observed and imaged at 0, 12, and 24 hours. Wound areas were measured using ImageJ software with the “Wound Healing Size Tool” plugin (*128*), and the measured wound areas at each time point were normalized to the 0-hour measurement.

Cell migration was evaluated using 8*μm* cell culture inserts for a 24-well plate (725301, NEST). After 24 hours of siRNA transfection, cells were digested, and 2 × 10^4^ cells suspended in 200*μL* of serum-free medium were seeded into the inserts. To attract migrating cells, 600*μL* of complete medium was added to the bottom wells. After 24 hours, non-migrating cells in the inserts were removed with cotton swabs. Inserts were fixed with 4% paraformaldehyde for 20 minutes, washed once with PBS, and stained with crystal violet (C0121, Beyotime) for 30 minutes. Migr125 ated cells were imaged under a microscope. A single image was taken per well, and the number of stained cells was quantified using the “Analyze Particles” function in ImageJ.

#### Kaplan-Meier survival analysis

Kaplan-Meier survival analysis was conducted using the Kaplan-Meier Plotter (KMplot) platform (https://kmplot.com). For THAP2, we focused on TNBC patients, defined by the absence of expression of ER, PR and HER2, and high expression of KI67 (a characteristic of TNBC). To ensure cohort specificity, we applied the following filters in KMplot: ER status = negative, PGR status = negative, HER2 status = negative, and KI67 status = positive. This yielded 77 RNA-seq samples. Since the RNA-seq data of OV patients is not available in KMplot, we analyzed gene-chip data from 1,435 samples. The overall survival plots were generated directly from the KMplot platform.

### Statistical tests

This study employed the one-sided Mann-Whitney U test to evaluate various datasets and assumptions.

We assessed the differences in SCCs between the PertFormer Attention Score and master-TF ChIP-seq signals (**Fig. 2A**). We tested whether the SCCs of the PertFormer attention scores for included cells, excluded cells, included genes, and excluded genes were significantly higher than the SCCs of scATAC-seq signals.

A differential analysis of the peak ranks, derived from PertFormer and ATAC-seq technologies for CRISPRi-FlowFISH and VISTA-validated EG pairs, was also conducted using the one-sided Mann-Whitney U test. This aimed to ascertain whether peak ranks of true EG pairs are significantly lower than those of false EG pairs (**Fig. 3A, B**). Similarly, predictions of gene expression changes by PertFormer, in response to *in silico* KO of enhancer and repressor loci, were statistically compared for up- and down-regulated genes using the same non-parametric test (**Fig. 3D**). For up-regulated genes (RG-pairs), gene expression differences (KO-Ref) were tested to see if they are significantly higher than zero, and for down-regulated genes (EG-pairs), if they are significantly lower than zero. In the K562 cell line, PertFormer’s gene expression predictions, following *in silico* TF perturbations, were examined using one-sided Mann-Whitney U tests (**Fig. 3F**). For up-regulated DEGs, the gene expression differences (Perturb-Ref) were tested to determine if they are significantly higher than zero, and for down-regulated DEGs, if they are significantly lower than zero.

To validate the effectiveness of PertFormer in identifying causal SNPs that affect gene expression, we applied the one-sided Mann-Whitney U test to verify if the percentile peak ranks of PertFormer and ATAC-seq for causal SNPs are significantly lower than those for non-causal SNPs within the GTEx V8 dataset (**Fig. 4A** and **Fig. S6A**). Moreover, The one-sided Mann-Whitney U test was utilized to assess if the attention score differences for causal SNPs, as measured by PertFormer-Elementary Transformer-1 between reference and alternate sequences, were significantly greater than those observed for non-causal SNPs within the same GTEx V8 dataset (**Fig. 4B**). Analyses also extended to the predicted expression changes after *in silico* ATAC signal deletions at SNP locations associated with up/down-regulation; these differences were statistically examined using a one-sided Mann-Whitney U test (**Fig. 4E**). For ATAC deletions at SNPs with up-regulatory effects, gene expression differences (KO-Ref) were tested to see if they are significantly lower than zero, and for deletions at SNPs with down-regulatory effects, if they are significantly higher than zero.

One-sided Mann-Whitney U test was similarly employed to compare the genome-wide influence, *in silico*OSKM KI to human fibroblast, human cardiomyocytes, and human B cells (**Fig. 5A**). For down-regulated DEGs, gene expression differences (KI-Ref) were tested to see if they are significantly higher than zero, and for up-regulated DEGs, if they are significantly lower than zero. For biological validation *in vitro*, all experiments were performed with at least three biological replicates. Quantitative data were processed with R (v4.3.3) and are presented as *mean*±*SEM*. The significance of differences between the target and control groups was assessed using an unpaired, two-tailed Student’s t-test.

## Supplementary figures

**Figure S1:**
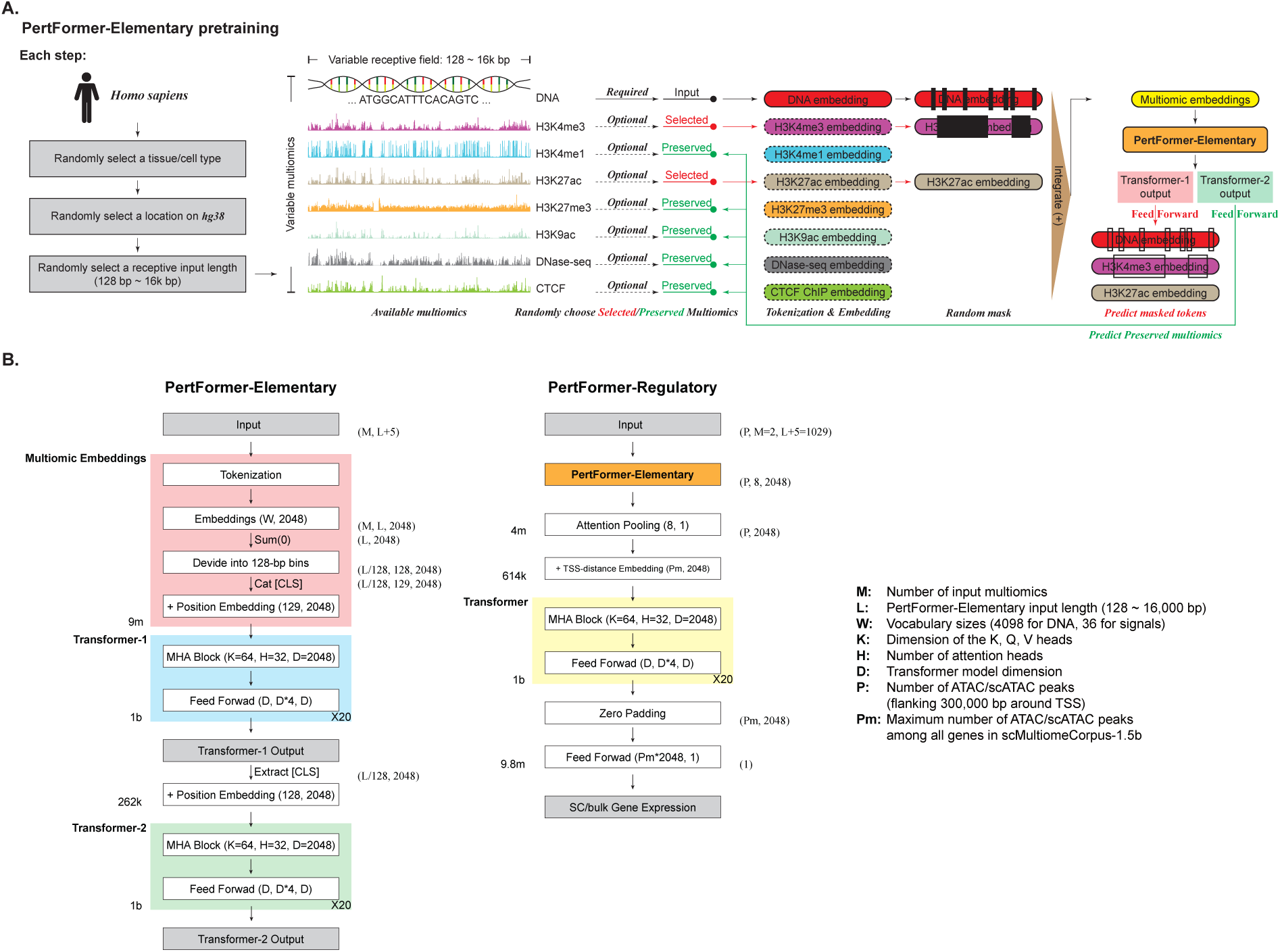
Details of PertFormer architecture and pretraining. **A.** Pretraining process of PertFormer-Elementary. In each step, the pretraining multiomic data of DNA sequence and 7 kinds of epigenetic signals were sampled from a random tissue or cell type with a random receptive length at a random location on the GRCh38 assembly, forming the bulkMultiomeCorpus-55b. Then, these epigenetic markers were randomly classified into Selected and Preserved groups. DNA sequence along with the signals from Selected group were randomly masked and input to PertFormer-Elementary. The output of Transformer-1 was used to reconstruct the masked tokens, and the output of Transformer-2 was used to predict the signals from the Preserved group. These two tasks were performed simultaneously. **B.** Detailed architectures of PertFormer model. Output shapes were showed as tuples on right side of each block. Annotations of hyper-parameters were presented on the right.

**Figure S2:**
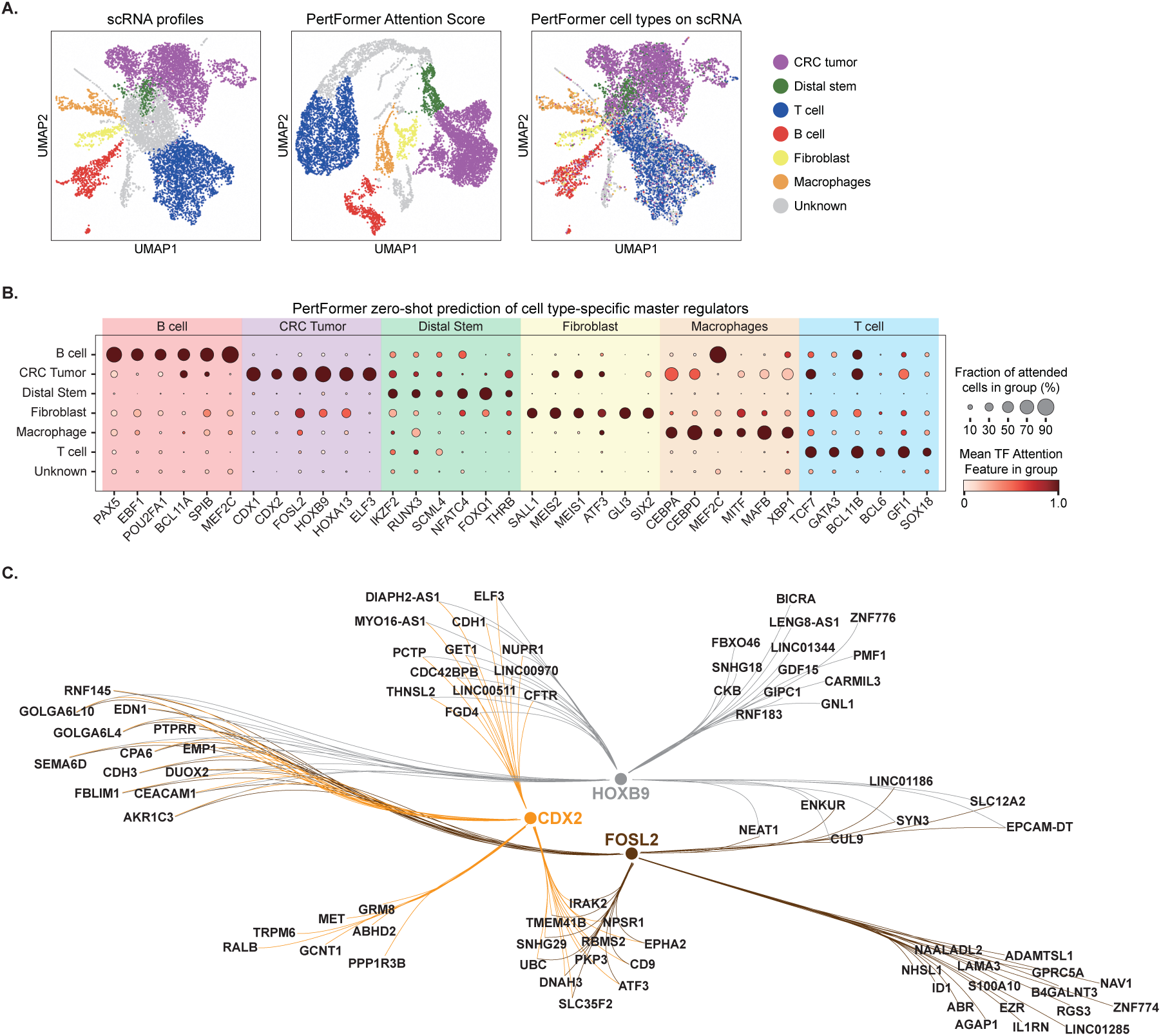
An Excluded-cell GRN inference case in CRC tumor tissue. This tissue was not included in scMultiomeCorpus-1.5b, and all predictions were generated without further training. **A.** UMAP of single cells from CRC tumor tissue based on experimental scRNA profiles (left) and PertFormer Attention Feature (middle). Cell types annotated by Attention Feature were marked on the UMAP of experimental scRNA (right). Cell type annotation results between scRNA and PertFormer showed a high consistency of 0.802 (right). **B.** Cell type-specific master regulators predicted by PertFormer Attention Feature. Mean Attention Features were shown by color scale, while fraction of cells were shown by size scale. Many well-known master regulators of each cell type were recovered by PertFormer. **C.** PertFormer-predicted GRNs regulated by CDX2, HOXB9, and FOSL2 in tumor cells of CRC. These 3 TFs formed 7 distinct groups of regulatory combinations.

**Figure S3:**
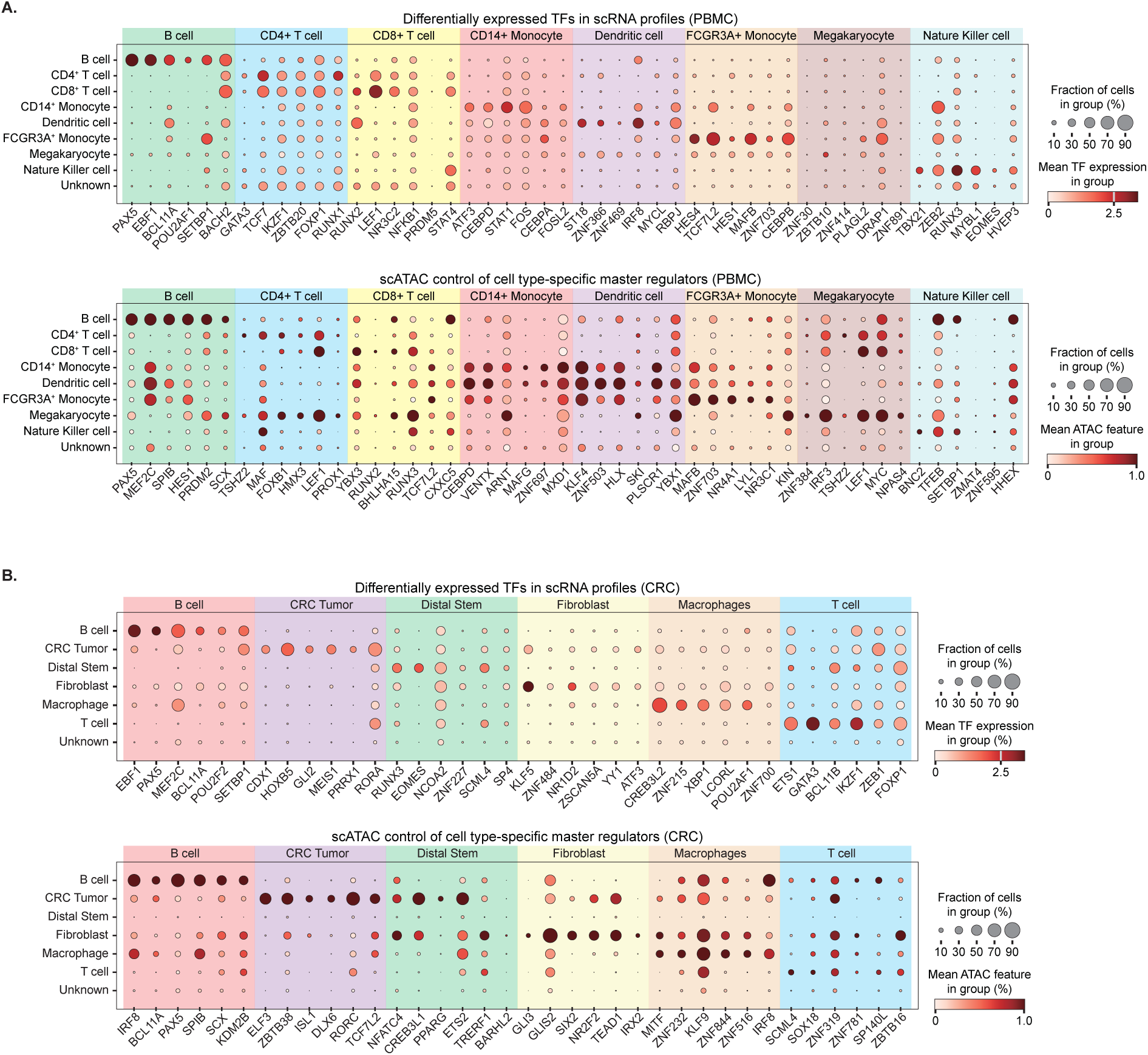
Control groups of master regulator predictions in PBMC and CRC. For scRNA profiles, differentially expressed TFs for each cell type were presented. For scATAC controls, the Attention Scores in **Equation**10 were replaced by scATAC signals. PertFormer-predicted master regulators showed substantial differential expressions. Those master regulators cannot be discovered by the raw scATAC profiles.

**Figure S4:**
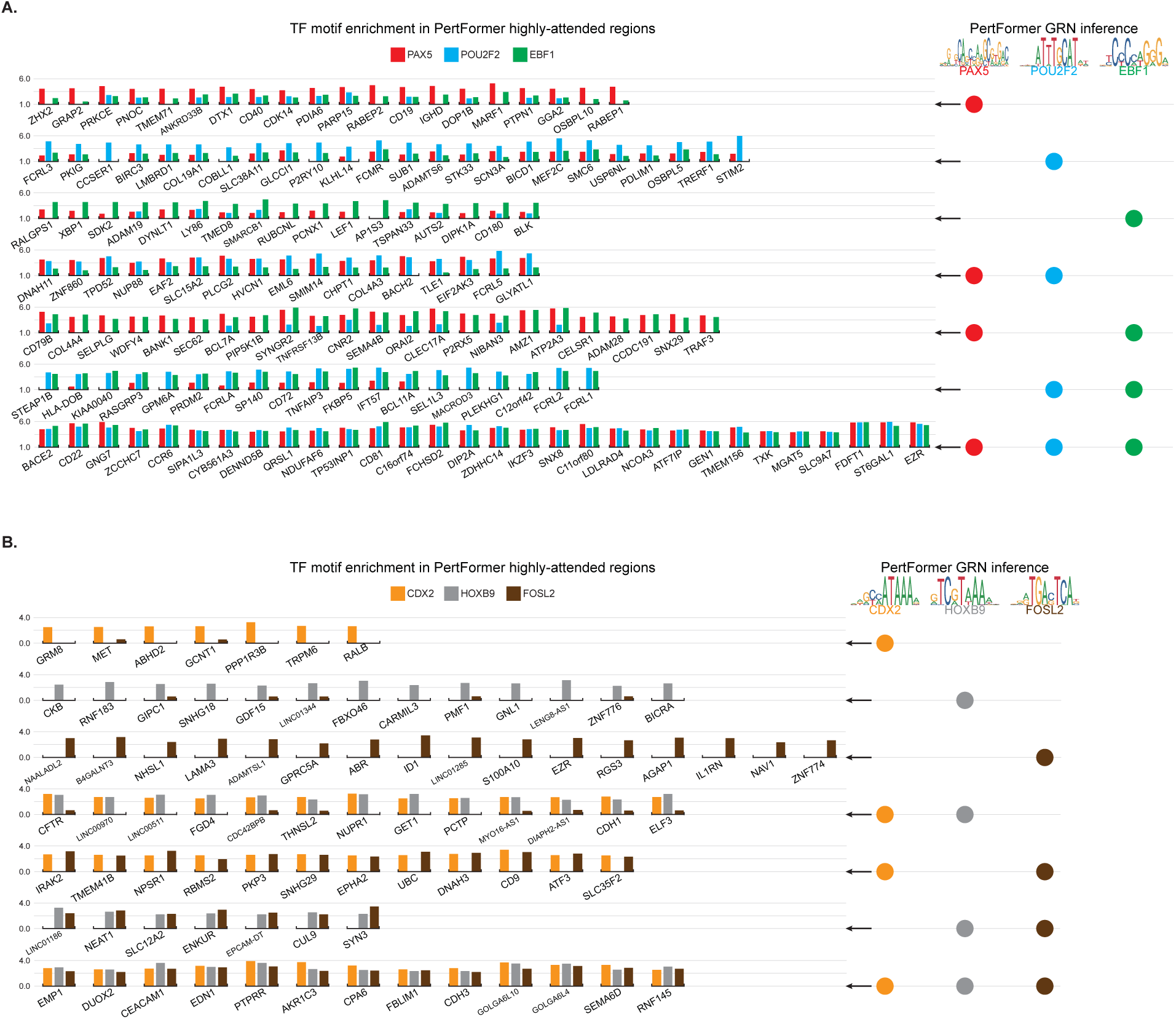
**A.** TF motif enrichment details in PertFormer highly-attended regions across different genes in B cells of PBMCs. Genes were classified into 7 groups by their enrichment motif score of TFs, forming PertFormer’s GRN inference in **Fig. 2C**. **B.** TF motif enrichment details in PertFormer highly-attended regions across different genes in tumor cells of CRC. Genes were classified into 7 groups by their enrichment motif score of TFs, forming PertFormer’s GRN inference in **Fig. S2**.

**Figure S5:**
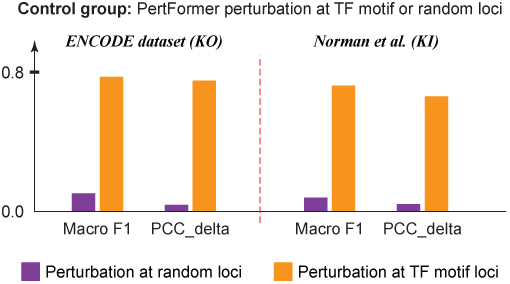
Control group to investigate the influence of KO and KI loci for TF perturbations. We conducted ATAC signal deletions (TF KO) or insertions (TF KI) at random loci for each gene. PertFormer lost predictive ability when the perturbations were applied without considering TF motifs (purple bars). These validated the methods of PertFormer for predicting TF perturbations by modifying the ATAC signals at TF motif loci.

**Figure S6:**
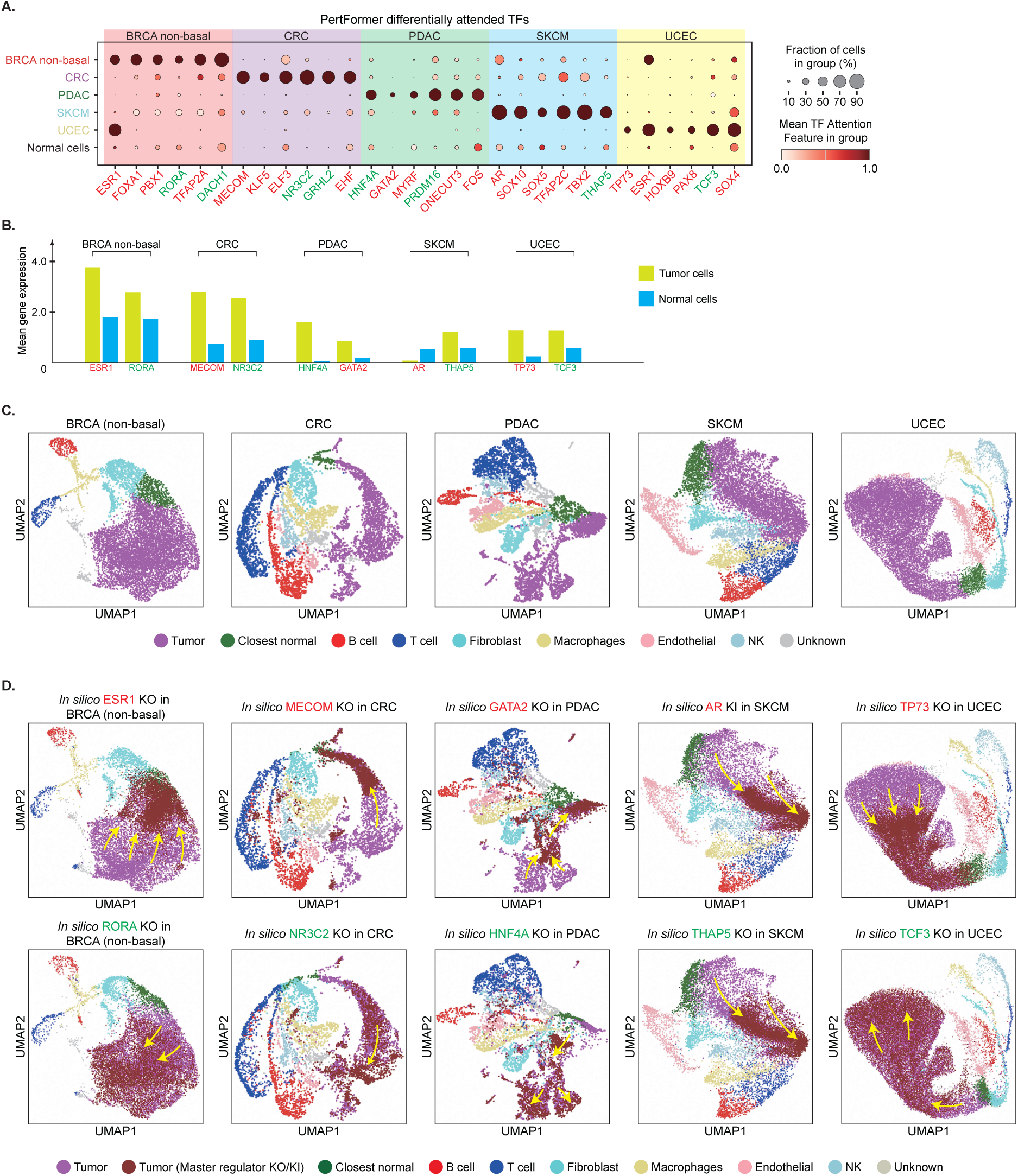
**A.** Predictions of master regulators in 5 tumor types. The top 6 predicted TFs in each tumor type were shown. For each tumor type, tumor-promoting TFs were marked in red, while tumor-suppressive TFs were marked in green. Mean TF expressions and fraction of cells were shown by color scale and size scale, respectively. **B.** Pseudobulk expression levels of the predicted TFs in tumor versus normal cells. **C.** The original UMAP of unperturbed normal and tumor single cells in each of the 5 tumor tissues, based on PertFormer Attention Feature. **D.** UMAP of normal, tumor, and *in silico* perturbed tumor cells in each of the 5 tumor tissues, based on PertFormer Attention Feature. For each tumor type, we respectively perturbed one tumor-promoting and one tumor-suppressive regulators which ranked on top of PertFormer’s predictions (the TFs with the most distinct Attention Features). KO/KI simulation was performed if the TF expression was higher/lower in tumor cells compared to normal cells. The *in silico* perturbations were performed on all tumor single cells (purple dots in each plot), and the perturbed cells were marked in brown dots. Here, tumor-suppressive effects of perturbation can be reflected as cell state transitions from tumor cells towards CNCs (KO of ESR1, MECOM, GATA2, and TP73). KI of AR, and KO of RORA, NR3C2, HNF4A, THAP5, and TCF3 showed tumor-promoting effects, as the perturbed cells were transited away from the CNCs. All the simulated effects corresponded with previous studies.

**Figure S7:**
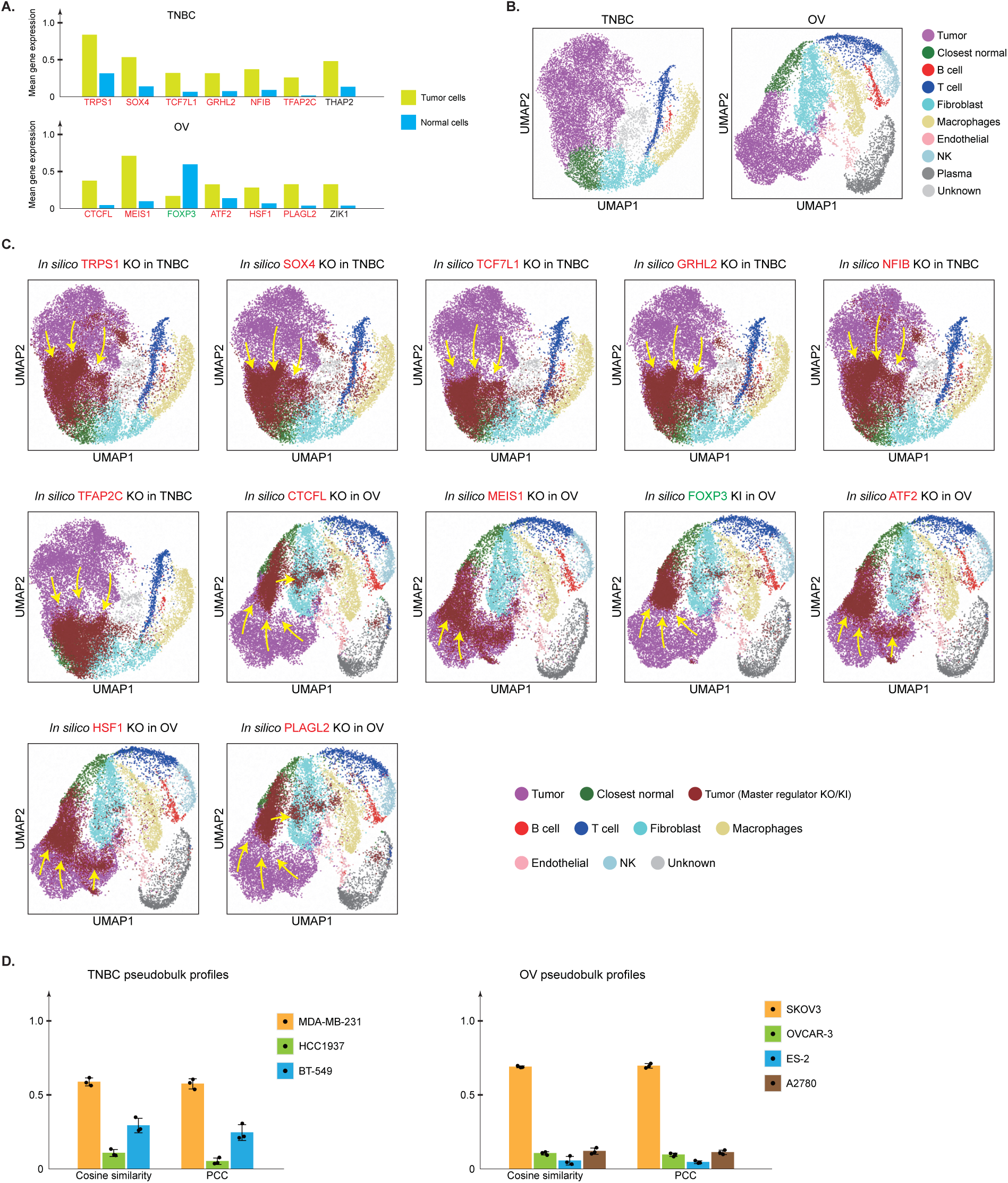
**A.** Pseudobulk expression levels of the predicted TFs in tumor versus normal cells. For each tumor type, previously-reported tumor-promoting TFs were marked in red, while previously-reported tumor-suppressive TFs were marked in green. **B.** The original UMAP of unperturbed normal and tumor single cells in TNBC and OV tumor tissues, based on PertFormer Attention Feature. **C.** UMAP of normal, tumor, and *in silico* perturbed tumor cells in TNBC and OV tumor tissues, based on PertFormer Attention Feature. In TNBC and OV tumors, perturbations were simulated respectively for each of the TFs ranked before the novel targets. Specifically, perturbation of TET3 was not performed as its motif was not available. Perturbation of USF3 was not simulated because its functions in OV were inconclusive. KO/KI simulation was performed if the TF expression was higher/lower in tumor cells compared to normal cells. The *in silico* perturbations were performed on all tumor single cells (purple dots in each plot), and the perturbed cells were marked in brown dots. Here, all the perturbations showed tumor-suppressive effects, which can be reflected as cell state transitions from tumor cells towards CNCs. These simulated effects corresponded with previous studies. **D.** Cosine similarity and PCC between pseudobulk expression profiles of tumor cells, and RNA-seq-measured expression profiles of different cancer cell lines. MDA-MB-231 and SKOV-3 were the closest cell line of the TNBC and OV tumor cells.

